# *In vivo* RUBISCO activity in *Synechocystis* is regulated by RuBP availability

**DOI:** 10.1101/2025.06.06.658252

**Authors:** Luisa Wittemeier, Stéphanie Arrivault, Niels Neumann, Boris Maček, Nils Schmidt, Martin Hagemann, Joachim Kopka

## Abstract

Ribulose-1,5-bisphosphate carboxylase/oxygenase (RUBISCO) is the main CO_2_-fixing enzyme on earth and entry point of carbon into the Calvin-Benson-Bassham cycle. Fueled by photosynthesis, C-assimilation by RUBISCO must be tightly controlled. RUBISCO regulation upon transition from light to darkness is not fully understood in the cyanobacterium *Synechocystis* sp. PCC 6803 (*Synechocystis*). *Synechocystis* does not have a RUBISCO activase that regulates RUBISCO activity in vascular plants by removing intrinsic sugar phosphate inhibitors from its active site. Instead, the regulatory CP12 protein of *Synechocystis* inactivates glyceraldehyde 3-phosphate dehydrogenase (GAPDH2) and phosphoribulokinase (PRK) during darkness. This mechanism indicates metabolic regulation of RUBISCO. We investigated C-assimilation *in vivo* at the transition to darkness by dynamic ^13^CO_2_ labeling experiments. We monitored RUBISCO activity by ^13^C-incorporation into 1-C position of 3PGA. Other than the wild type, the *Δcp12* mutant continued to assimilate ^13^CO_2_ into 3PGA in darkness. RUBISCO abundances and specific activities were not altered in *Δcp12* and upon light to dark transition. CP12 was required to shut down the CBB cycle during the night. Complementation of *Δcp12* by native CP12 (*Δcp12::cp12*) and CP12 with mutated conserved cysteines in its GAPDH2- and PRK-binding domains (*Δcp12::cp12ΔCys*) showed that both native binding domains are required to fully inactivate the CBB cycle in the night. RuBP levels were highly elevated in *Δcp12* upon transition to darkness. Complementation with mutated and native CP12 variants gradually reduced RuBP to wild type levels and revealed highly significant correlation between RuBP concentration and the time-shifted ^13^C-uptake into 3PGA. We propose that RUBISCO activity in *Synechocystis* at day-night transition is regulated through depletion and blocked regeneration of RuBP. ^13^C-positional analyses of aspartate suggest regeneration of RuBP in *Δcp12* via dysregulated gluconeogenesis and the oxidative pentose phosphate path. We demonstrate that RUBISCO activity of *Synechocystis* is present throughout diurnal growth and depends on the availability of its substrate.

## Introduction

The ancient cyanobacteria were the first organisms doing oxygenic photosynthesis and adding oxygen to the atmosphere (Dismukes et al., 2001). They are believed to be the ancestor of the chloroplast in algae and higher plants taken up by a eukaryotic cell through endosymbiosis (Margulis, 1970). Recently, cyanobacteria gained application in green biotechnology, producing valuable substances based on light energy (Hagemann and Hess, 2018).

Cyanobacteria adapted over billions of years to live under diurnal light changes. To adjust cellular metabolism to continuous diurnal light and dark phases, they evolved a circadian clock (Köbler et al., 2024; Berwanger et al., 2025). During the day, cyanobacteria perform an autotrophic life style using photosynthesis. The captured light energy is converted to ATP and NADPH which fuel the Calvin-Benson-Bassham (CBB) cycle and CO_2_ fixation by ribulose-1,5-bisphosphate carboxylase/oxygenase (RUBISCO). During darkness, they must switch their metabolism to a heterotrophic-like mode, in which catabolism of stored glycogen is active to sustain viability. Glycogen catabolism in cyanobacteria is mainly via oxidative pentose phosphate (OPP) pathway, which also employs some enzymes of the CBB cycle in catabolic direction (Yang et al., 2002; Lucius and Hagemann, 2024). The changes between autotrophy and heterotrophy involve a lot of fundamental regulatory steps which still need to be investigated. The lack of compartmentalization in cyanobacteria requires tight control of metabolic reactions, especially at the protein and metabolite levels. Some strains such as the model *Synechocystis sp.* PCC 6803 (hereafter *Synechocystis*) can also grow with glucose as organic carbon source, i.e. this strain permits to compare autotrophic (light with CO_2_), mixotrophic (light with CO_2_ and glucose) and heterotrophic (dark with glucose) growth modes.

RUBISCOs paired with oxygenic photosynthesis produce the vast majority of organic carbon in the biosphere (Bar-On and Milo, 2019). RUBISCO as the key enzyme of the CBB cycle fixes CO_2_ in ribulose-1,5-bisphosphate (RuBP) resulting in 2 molecules 3-phosphoglycerate (3PGA). Due to its affinity to oxygen in the present day oxygen-rich atmosphere, RUBISCO has a wasteful oxygenase reaction producing 2-phosphoglycolate (2PG) and inhibiting carbon metabolism (Cleland et al., 1998). To avoid this reaction, different inorganic carbon concentrating mechanisms (CCMs) evolved, supporting the accumulation of CO_2_ around RUBISCO (Raven et al., 2008). The cyanobacterial CCMs consist of inorganic carbon uptake systems that actively accumulate HCO_3_^−^ inside the cell and carboxysomes, a prokaryotic micro-compartment that encapsulate RUBISCO and carbonic anhydrase to convert bicarbonate into CO_2_ (Price et al., 2008; Hagemann et al., 2021).

RUBISCO activity is usually light-dependent, because the ATP to produce its substrate RuBP and NADPH to reduce the initial fixation product 3PGA in the CBB cycle are provided by the photosynthetic complexes capturing light energy. A functional competent RUBISCO needs to be activated by carbamylation, which leads to an active RuBP binding site (Lorimer et al., 1976). Carbamylation was found to be regulated by binding of orthophosphates in *Synechocystis* (Marcus and Gurevitz, 2000). In plants, RUBISCO activity is further regulated by inhibition of sugar phosphates (as for example RuBP itself or RUBISCO by-products) and the removal of sugar phosphates by RUBISCO activase (Portis et al., 1986). Even though there are cyanobacteria with RUBISCO activase, e.g. *Nostoc* sp. PCC 7120 (Flecken et al., 2020), the regulation of RUBISCO activity in *Synechocystis* is still unclear, because a RUBISCO activating protein is absent from the genome (Lechno-Yossef et al., 2020). Correspondingly, it has been shown that *Synechocystis* RUBISCO is insensitive to different highly phosphorylated sugar inhibitors (Pearce, 2006).

To allow continuous substrate availability for RUBISCO during the day, RuBP is regenerated from 3PGA via the CBB cycle. Hence, the entire CBB cycle must be activated in the early morning. One main regulatory protein of the CBB cycle is the CP12 protein (Wedel and Soll, 1998), which activates and inactivates GAPDH2 and PRK in light/dark cycles in a redox-dependent manner in *Synechocystis* (Blanc-Garin et al., 2022; Lucius et al., 2022). Two different GAPDHs are encoded in the genome of *Synechocystis.* GAPDH1 is involved in glycolytic sugar catabolism and uses NAD(H), whereas GAPDH2 is part of the CBB cycle and uses NAD(H) and NADP(H) (Koksharova et al., 1998; Rajarathinam et al., 2025). Canonical CP12 proteins contain two cysteine pairs, one of which is located at the C-terminus and the second at the N-terminus (McFarlane et al., 2019). CP12 proteins are redox-regulated, mainly through thioredoxins but also through other physiological oxidants as H_2_O_2_, oxidized DTT, GSSG and GSNO (Marri et al., 2014). In darkness, the complex is formed through initial binding of two oxidized CP12 to each of two GAPDH tetramers via disulfide bonds at the C-terminal cysteines which is obligatory to build the complex with PRK. GAPDH needs to be loaded with NAD(H) instead of NADP(H), which is less abundant under oxidizing conditions (high NAD(H)/NADP(H) ratio; Tamoi et al., 2005). Two PRK dimers are recruited to each of the two intermediate GAPDH-CP12 complex at the N-terminal cysteines (Gurrieri et al., 2021). The complex formation leads to inactivation of GAPDH2, which catalyzes production of glyceraldehyde-3P under reduction of NAD(P)H, and PRK, which is necessary for RuBP synthesis under ATP consumption. Thus, both enzymes would dissipate a lot of energy equivalents if not inactivated during the night. In addition to the cysteine residues, the conserved negatively charged AWD_VEEL motif was found to be involved in the inactivation of PRK activity by interaction with positive residues of PRK (Gurrieri et al., 2021), and in regulating the redox equilibrium of NADPH (Blanc-Garin et al., 2022). Growth experiments with a *Synechocystis* mutant lacking CP12 (*Δcp12*) showed that it is dispensable for photoautotrophic growth and no growth effect was seen in light/dark cycles, whereas growth is strongly affected when glucose is added to the medium (Blanc-Garin et al., 2022; Lucius et al., 2022). CP12 proteins are universally distributed among all oxygenic photosynthetic organisms (except the parsinophyte Osterococcus; Groben et al., 2010; Lopez-Calcagno et al., 2014). In *Arabidopsis thaliana* and other angiosperms, three CP12 proteins were found, with CP12-1, and CP12-2 being highly homologous and CP12-3 rather distinct (López-Calcagno et al., 2017). The different versions showed redundancy in their functions. Knockout mutants of all CP12 versions lead to a decrease of PRK protein abundance and lower photosynthetic activity. In gymnosperms and *Chlamydomonas reinhardtii*, only one CP12 protein was found. All CP12 proteins have conserved N- and C-terminal cysteine pairs and AWD_VEEL motifs, which are used for the regulation of CBB cycle activity through inactivation of PRK and GAPDH. Additionally, there are CP12-like proteins in some cyanobacteria that can be fused to CBS (cystathionine β-synthase) domains, and that are only binding and inactivating PRK but not GAPDH (Stanley et al., 2013).

So far it is unknown how RUBISCO activity is regulated at day night transition in *Synechocystis*. We investigated the RUBISCO reaction and parts of the central carbon metabolism with dynamic ^13^CO_2_ labeling experiments at day night transition in WT and *Δcp12* to show that RUBISCO activity is regulated on a metabolic level by substrate availability. We used ^13^C positional enrichment analysis of 3PGA to investigate RUBISCO activity *in vivo*. The studies were complemented by proteomics and specific RUBISCO *in vitro* activity measurements that showed no differences between *Δcp12* and WT. Different *Δcp12* complementation mutants lacking the N- and C-terminal cysteine binding domains were included. These mutants showed intermediate RUBISCO activity, which strongly correlated with RuBP availability in a time-shifted manner. We concluded that RUBISCO activity in the night depends on substrate availability, with CO_2_ being available belated due to down-regulated CCM during darkness and RuBP only being available when PRK and GAPDH2 are active. ^13^C positional enrichment analysis of aspartate revealed that regeneration of RuBP during the night in *Δcp12* is via gluconeogenesis and upper OPP pathway, which is a futile cycle with no benefits but loss of ATP.

## Results

### RUBISCO activity in *Δcp12* continues at the beginning of the night

Changes in metabolite pools in the cell due to changes in environmental factors can indicate differences in enzyme activity, either of producing or of degrading enzymes. When transferring cells from day to night conditions, 3PGA concentration in *Synechocystis* WT cells was decreasing likely due to reduced RUBISCO activity during darkness (**Figure 1 A**). A mutant lacking the regulatory protein CP12, namely *Δcp12*, initially showed similar behavior as WT but strongly accumulates 3PGA after 60-90 min to 2.85 nmol*OD_750_^−1^*mL^−1^ (**Figure 1 A+B**). Additionally, RUBISCO oxygenase product 2PG is accumulating more than 15-fold in *Δcp12* compared to WT and *Δcp12::cp12* after shifting to darkness to 0.49 nmol*OD_750_^−1^*mL^−1^ (**Figure S 1 B**). This leads to the assumption that RUBISCO was active in general in *Δcp12* with focus on oxygenase activity at beginning of the night and transition to carboxylase activity after 60-90 min. To test this hypothesis, we subjected the cells to stable isotope labeled ^13^CO_2_ at beginning of the night. Indeed, ^13^CO_2_ was incorporated into 3PGA in *Δcp12* but not in WT and *Δcp12::cp12* (**Figure S 1 E**). Downstream metabolites 2PGA and PEP were enriched in ^13^C to a similar proportion as 3PGA in *Δcp12* (**Figure S 1 G-H**). Resolution of single carbon positions of the 3PGA backbone according to Rajarathinam et al. (2025) allows to specify that the main ^13^C enrichment was assimilated in 3PGA 1-C (**Figure 1 C**), which is directly incorporated through RUBISCO. After 60-90 min, ^13^C was also found in 3PGA 2,3-C (**Figure 1 D**), which only gets label through recycling by CBB cycle (Rajarathinam et al., 2025). This is confirmed by 2PG which as well got label after 60-90 min (**Figure S 1 F**). The accumulation of 2PG in *Δcp12* after transition to darkness could not be prevented by removing O_2_ from the fresh culture medium by N_2_-bubbling indicating that there were internal O_2_ pools leading to preferred oxygenase activity (**Figure S 2 C**).

**Figure 1.**
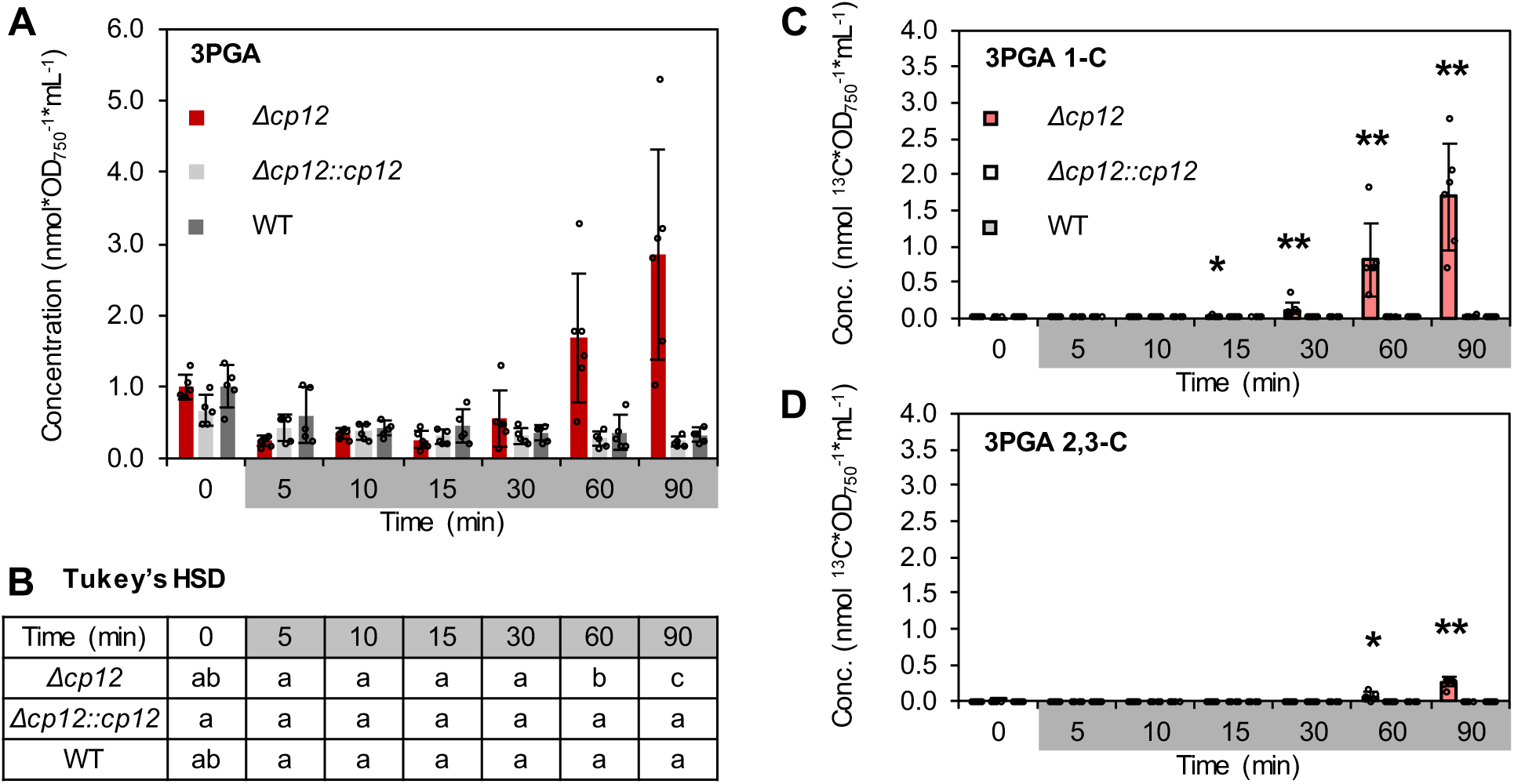
RUBISCO activity *in vivo* in *Δcp12* during darkness. *Synechocystis Δcp12* was cultivated in comparison to *Δcp12::cp12* and WT. Sampling and ^13^CO_2_ labeling was done at transition to the fourth night. 3PGA levels were determined using GC-EI-MS, ^13^C enrichment using GC-APCI-MS. **A** 3PGA concentration (nmol*OD_750_*mL^−1^) in *Δcp12*, *Δcp12::cp12* and WT. **C+D** E^13^C concentration (nmol*OD_750_*mL^−1^) of 3PGA 1-C (**C**) and 2,3-C (**D**). Bars show the average of 5 independent biological replicates ± standard deviation. Asterisks show significant differences between *Δcp12* and WT and *Δcp12::cp12* (tested independently) based on Wilcoxon-Mann-Whitney testing (*: *P*<0.05, **: *P*<0.01). Letters in the table (**B**) display significant differences of 3PGA concentration between all conditions based on Tukey’s HSD (*P*<0.05).

To determine whether deletion of *cp12* also enables CO_2_ assimilation by RUBISCO at the end of the night, we labeled and sampled *Synechocystis* WT, *Δcp12* and *Δcp12::cp12* at the end of the night, starting the labeling pulse 90 min before transition to day conditions. 3PGA and 2PG concentrations were still increased in *Δcp12* compared to WT and *Δcp12::cp12* (**Figure S 3 A-D**). E^13^C could be found after 60-90 min in 1,2,3-C and 1-C of 3PGA in *Δcp12*, but not in 2,3-C of 3PGA, indicating that RUBISCO is still active but regeneration of RuBP via CBB cycle strongly reduced (**Figure S 3 E-G**). Hence, CP12 is needed throughout the night to inactivate GAPDH2 and PRK and preventing RUBISCO activity.

Since RUBISCO seems to be in an active state in *Δcp12* at the end of the night, we checked if there is an advantage at beginning of the next day. However, no higher initial RUBISCO activity could be found, since ^13^C concentration in 1-C did not differ between the strains in the early time points (**Figure S 4 H**). Elevated 3PGA levels at the end of darkness directly aligned with WT levels upon turning on the light (**Figure S 4 A**). Minor differences could be observed in the long-term (60-90 min) regeneration rate. *Δcp12* showed an increased amount of ^13^C in 2,3-C of 3PGA compared to WT and *Δcp12::cp12* (**Figure S 4 F+I**).

Overall, CP12 had the strongest effect at beginning of the night when it inactivates GAPDH2 and PRK to shut down CBB cycle and consequently prevents CBB cycle and thereby RUBISCO activity. Hence, we further investigated day/night transition and the influence of CP12 on RUBISCO activity during the night.

### No regulation of RUBISCO abundance and specific activity at beginning of the night

Since cyanobacteria are prokaryotic organisms, they lack compartmentalization. Consequently, a lot of regulatory steps are at protein and metabolite level. We checked RUBISCO protein abundance by detecting RUBISCO large subunit (RbcL) with Western Blot (**Figure 2 A**). There were no measurable differences between WT, *Δcp12::cp12* and *Δcp12*, neither at the end of the light period, nor at the end of our labeling period 90 min after shift to darkness. The same conditions were tested with proteomics. There were no significant changes in RbcL, RUBISCO small subunit (RbcS), or RUBISCO chaperone (RbcX) that could explain higher RUBISCO activity during the night in *Δcp12*. We also checked the effect of *Δcp12* on all proteins involved in central carbon metabolism (**Figure S 5 A**). The log_2_-fold changes between conditions were between 1 and −1, meaning that none of the proteins changed more than a factor of 2. Only few changes were significant but none would explain higher CO_2_ assimilation during the night in *Δcp12*. The only protein that strongly accumulated in *Δcp12* in comparison to WT and *Δcp12::cp12* was PhaP (**Figure S 5 C**), known to regulate the size of PHB granula and potentially PHB synthase activity (Hauf et al., 2015). PHB related proteins were seen before to accumulate upon cellular stress due to unbalanced nutrient and carbon conditions (Koch et al., 2020).

**Figure 2.**
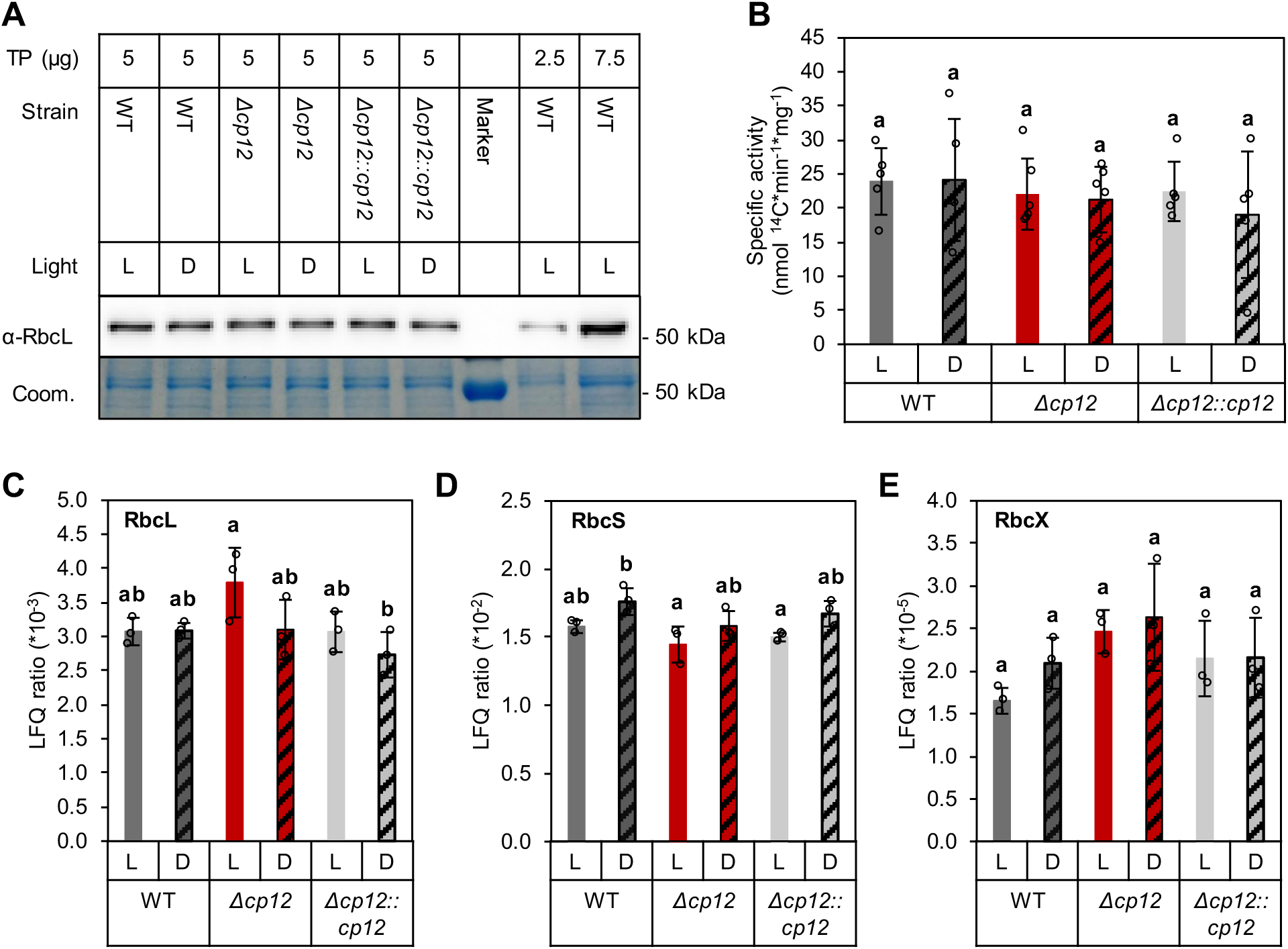
RUBISCO activity is not regulated on protein level. **A** RbcL detection with Western Blot using RbcL-antibody (α-RbcL) in *Δcp12*, *Δcp12::cp12*, and WT in light (L, t_0_) and darkness (D, 90 min after transition to night; TP :total protein, Coom: Coomassie brilliant blue). **B** Specific RUBISCO carboxylase activity was determined *in vitro* of the same conditions. **C-E** Proteomics was used to differentiate changes in RUBISCO large (**C**) and small (**D**) subunit and RUBISCO chaperon (**E**). Displayed are LFQ intensities normalized to the sum of LFQ intensities per sample (LFQ ratio). Letters in the diagramms display significance based on Tukey’s HSD (*P*<0.05).

To check whether the RUBISCO protein has different specific activities during the night, we measured RUBISCO *in vitro* activity with a ^14^C based radiometric measurement. There were no differences in specific activity between *Δcp12* and WT or *Δcp12::cp12*, neither during the day nor during the night (**Figure 2 B**). RUBISCO is not in a more active state in *Δcp12* during darkness, excluding that RUBISCO is regulated on enzyme level.

### Contribution of GAPDH2 and PRK activity to RUBISCO activity in the night

CP12 inactivates GAPDH2 and PRK via covalent binding through disulfide bonds. The cysteines facilitating GAPDH2 binding are located at the C-terminus, the cysteines for PRK binding at the N-terminus of the CP12 protein. *Δcp12* was complemented with *cp12* variants of which the cysteines were replaced by serines, either the C-terminal pair (*Δcp12::cp12ΔCysC*) or the N-terminal pair (*Δcp12::cp12ΔCysN*), or both C- and N-terminal cysteines (*Δcp12::cp12ΔCysNC*) (Lucius et al., 2022). The concentration of 3PGA dropped in all strains upon transition to darkness (**Figure 3 A+C**). The complete knockout strain *Δcp12* accumulated 3PGA after 60-90 min as seen before (**Figure 1 A**). All complementation strains showed WT behavior over the whole time course. However, RUBISCO oxygenase product 2PG increased directly after onset of the night strongly in *Δcp12*, but also in *Δcp12::cp12ΔCysNC* and to a lower level and not significant in *Δcp12::cp12ΔCysC*, indicating that RUBISCO activity in early time points is also higher if at least GAPDH2 binding is prohibited (**Figure 3B+D)**. The concentration of ^13^C in 3PGA 1,2,3-C was significantly increased in *Δcp12::cp12ΔCysC* after 30 min and in *Δcp12* and *Δcp12::cp12ΔCysNC* after 60 min (**Figure 3 E**). The absolute amount was highest in *Δcp12* due to higher concentration of 3PGA in the cells. Comparison of E^13^C shows that a similar proportion was labeled in *Δcp12* and *Δcp12::cp12ΔCysNC* and a smaller part in *Δcp12::cp12ΔCysC* (**Figure 3 H**). While *Δcp12::cp12ΔCysC* only showed ^13^C label in 3PGA 1-C, *Δcp12* and *Δcp12::cp12ΔCysNC* also got label in 3PGA 2,3-C after 60-90 min (**Figure 3 F, G, I, J**) indicating that regeneration is only possible if binding of both, PRK and GAPDH2, is prohibited. Indication of RUBISCO activity was highest when *cp12* or both cysteine pairs were deleted, meaning that lack of binding both enzymes is necessary for high assimilation rates. However, if only PRK binding through cysteines is prohibited, no effect on RUBISCO activity could be seen, whereas deletion of the GAPDH2 binding domains leads to minor activity.

**Figure 3.**
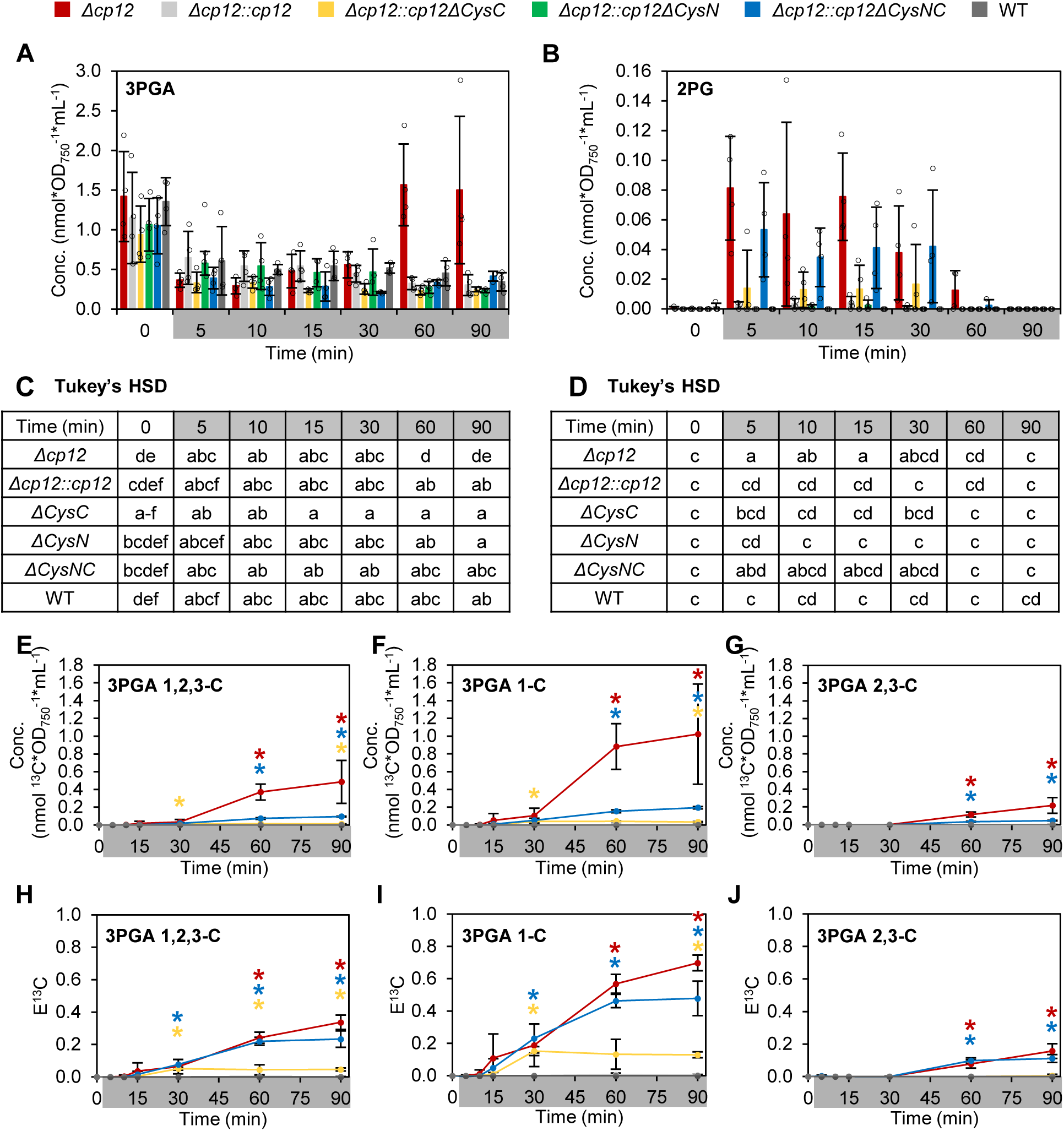
CO_2_ assimilation by RUBISCO and regenerative CBB cycle activity is dependent on available regeneration routes. *Δcp12* was complemented with variants of *cp12* lacking either one or both cysteine residues facilitating GAPDH2 and PRK binding (CysC: C-terminal cysteine, facilitates GAPDH2 binding; CysN: N-terminal cysteine, facilitates PRK binding; CysNC: N- and C-terminal cysteine). All strains were labeled and sampled at/after transition to the fourth night. **A + B** Changes in 3PGA and 2PG concentration (nmol*OD_750_*mL^−1^) at transition to the night. Letters in the tables (**C+D**) display significant differences of 3PGA (**C**) and 2PG (**D**) concentration between all conditions based on Tukey’s HSD (p<0.05). **E-G** Absolute concentration of assimilated ^13^C (nmol*OD_750_*mL^−1^) in 3PGA 1,2,3-C, 1-C and 2,3-C. **H-J** Relative ^13^C enrichment (E^13^C) of 3PGA 1,2,3-C, 1-C and 2,3-C. Asterisks are colored in the same way as data points and display significant differences to WT and *Δcp12::cp12* (tested independently) based on Wilcoxon-Mann-Whitney testing (*: p<0.05).

### RUBISCO activity is regulated by substrate availability

An enzyme can only catalyze its reaction if the substrate is provided in sufficient amounts. RUBISCO depends on two substrates: Ribulose-1,5-bisphosphate (RuBP) and CO_2_, standing in competition with the non-desired substrate O_2_. Since we find labeling of 3PGA in *Δcp12, Δcp12::cp12ΔCysNC* and *Δcp12::cp12ΔCysC*, RuBP and CO_2_ should be present during darkness in these strains. The accumulation of 2PG in *Δcp12, Δcp12::cp12ΔCysNC* and *Δcp12::cp12ΔCysC* after 5 min indicates that RuBP is already available directly after onset of the night, and that CO_2_ is lacking at the beginning of the night. This is probably an artefact, because all dissolved Ci was removed from the medium before the ^13^CO_2_ labeling pulse to enable rectangular labeling. We determined RuBP with LC-MS/MS in *Δcp12*, WT and all complementation strains. RuBP indeed accumulated strongly directly after transition to darkness up to 30 min in *Δcp12* to ∼2 nmol and a little less in *Δcp12::cp12ΔCysNC* (∼0.7 nmol) and *Δcp12::cp12ΔCysC* (∼0.6 nmol; **Figure S 6**, **Figure 4 A**). We determined ^13^C label in 3PGA 1,2,3-C in the same experiment. The relative concentration of ^13^C in 3PGA increased over time to ∼29.9% (maximum-scaled intensity multiplied with E^13^C) at 90 min in *Δcp12* (**Figure 4 B**). *Δcp12::cp12ΔCysNC* reached a relative ^13^C concentration of ∼4.1% and *Δcp12::cp12ΔCysC* reached ∼2.9%. The ^13^C enrichment in 3PGA occurs in a time-shift to the accumulation of RuBP (**Figure S 6**). We log_10_-transformed the data for RuBP concentration at 15 min and for 3PGA relative ^13^C concentration at 90 min to value small differences with the same intensity as big differences, as there is a huge range for the different strains in both, RuBP concentration and 3PGA relative ^13^C concentration. Correlation analysis (based on Pearson’s correlation) shows a very strong (*r* = 0.9040) and highly significant (*P* = 1.398*10^−9^) time-shifted correlation between RuBP concentration at 15 min and 3PGA relative ^13^C concentration at 90 min. The time-shifted correlation becomes obvious by correlating RuBP concentration at 15 min with 3PGA relative ^13^C concentration at 15-90 min (**Figure S 6**). There is already a significant correlation at 15 min, but significance and correlation coefficient increase until 60-90 min. RUBISCO activity during the night strongly depends on RuBP availability. Accumulation of RuBP during the night is prohibited in *Synechocystis* WT through inactivation of GAPDH2 and PRK by CP12. These results allowed us to conclude that RUBISCO carboxylase activity in *Synechocystis* is solely regulated by its substrates, RuBP and CO_2_.

**Figure 4.**
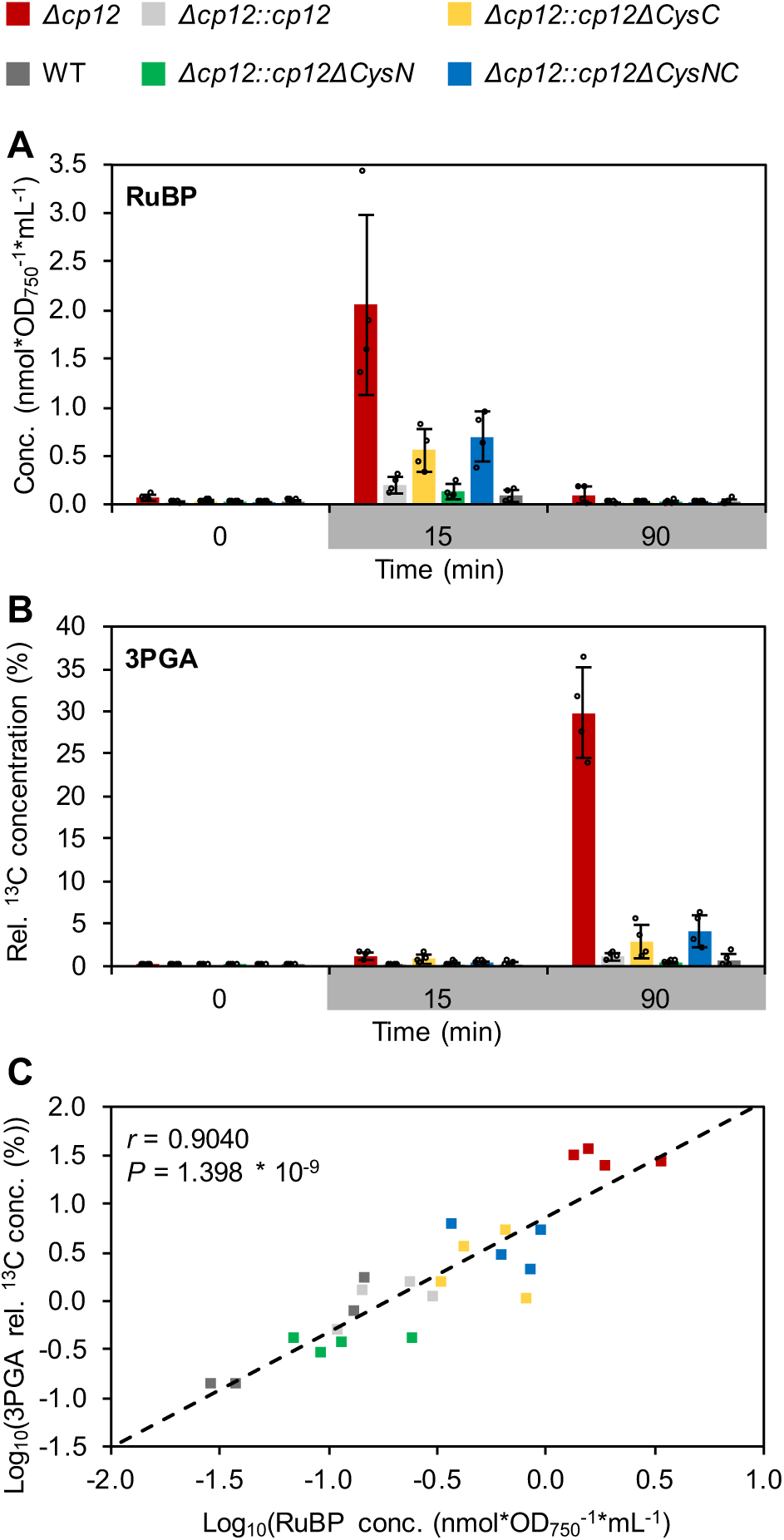
E^13^C in 3PGA correlates with RuBP availability in *Synechocystis* in the dark. *Synechocystis* WT and mutant strains were sampled and labeled with ^13^CO_2_ at transition to the night. **A** RuBP concentration (relative intensity, maximum-scaled) was determined using LC-MS. **B** 3PGA concentration and E^13^C were determined in the same LC-MS chromatograms and combined to E^13^C concentration (max-scaled 3PGA concentration multiplied with E^13^C fraction). **C** Log_10_-transformed 3PGA E^13^C concentration 90 min after transition to darkness is correlated with the log_10_-transformed RuBP concentration at 15 min with a correlation coefficient of 0.9040 (pearson correlation) and p-value of 1.398*10^−9^. Bar plots represent the mean ± SD of 4 independent biological replicates.

### Regeneration of RuBP via gluconeogenesis and OPP during the night

In *Synechocystis* (and other phototrophic organisms), RuBP regeneration via CBB cycle is shut down during the night. One main regulator is the redox regulated CP12 protein which inactivates PRK and GAPDH2 via covalent binding. In *Δcp12*, RuBP accumulated after transition to the night. This indicates that there is a potential of RuBP regeneration in this mutant. To get evidence which route is used, we have analyzed the positional labelling pattern of aspartate. Aspartate originates from oxaloacetate, which is generated by PEPC from phosphoenolpyruvate (PEP) and HCO_3_^−^. The carbon organization of 3PGA is kept in PEP and transferred to 1,2,3-C of aspartate. 4-C of aspartate originates from the assimilated carbon by PEPC (**Figure S 7**). In WT and *Δcp12::cp12*, only 4-C of aspartate is ^13^C enriched due to PEPC activity in the night (**Figure 5 C**). However, *Δcp12* also showed E^13^C in 1-C of aspartate after 60 min and in 2-C of aspartate after 90 min. No E^13^C could be detected in 3-C of aspartate (**Figure 5 C**).

**Figure 5.**
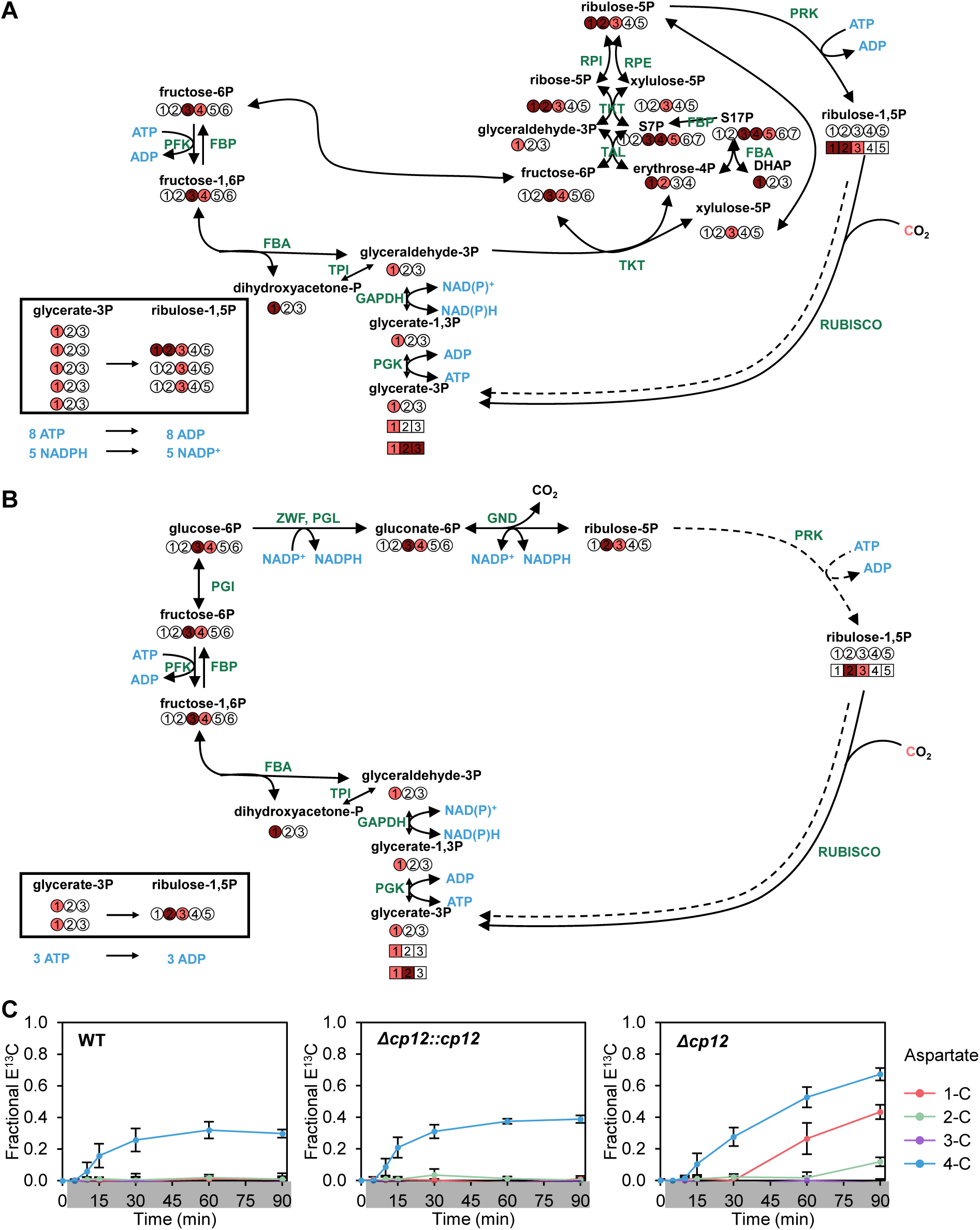
Regeneration of RuBP during the night in *Δcp12*. **A + B** Potential regeneration pathways of RuBP. Solid lines and circles indicate the labeling status of carbons representing the first round of RUBISCO assimilation, dashed lines and squared carbon labels a second round of RUBISCO assimilation. Light red carbon label is added by RUBISCO and transferred to glyceraldehyde-3P, whereas dark red label is further shuffled to dihydroxyacetone-P. **A** If RuBP is regenerated via CBB cycle, potential label at 1-C of 3PGA is shuffled to label in 1-C and 2-C to equal proportions. All regenerated RuBP molecules are labeled at 3-C. Three RuBP molecules with 1x label at 1-C, 1x label at 2-C and 3x label at 3-C are regenerated from 5 molecules 3PGA labelled at 1-C. **B** When RuBP is regenerated via gluconeogenesis and parts of the OPP pathway, 2 molecules 3PGA labeled at 1-C are combined to 1 molecule RuBP having the label at 2-C and 3-C. In the second round of RUBISCO assimilation, 3PGA molecules are either labeled at 1-C or at both, 1-C and 2-C, but not at 3-C. **C** Positional E^13^C in aspartate was determined in WT, *Δcp12* and *Δcp12::cp12* after transition to darkness (n=5, mean ± standard deviation). Difference in 2-C and 3-C labelling in *Δcp12* after 90 min was tested with Wilcoxon signed-rank test (*P* = 0.03125). Pathway modified based on Makowka et al. (2020).Enzyme abbreviations: PGK – Phosphoglycerate kinase, TPI – Triose-phosphate isomerase, FBA – Fructose-bisphosphate aldolase, FBP – Fructose-1,6-bisphosphatase, PFK – Phosphofructokinase, TKT – Transketolase, TAL – Transaldolase, RPI – Ribose-5-phosphate isomerase, RPE – Phosphopentose epimerase, PGI – Phosphoglucose isomerase, ZWF – Glucose-6-phosphate dehydrogenase, PGL – 6-Phosphogluconolactonase, GND – 6-Phosphogluconate dehydrogenase

To regenerate RuBP, 1-^13^C labelled 3PGA is stepwise converted to glyceraldehyde-3P (GA3P) and dihydoxyacetone-P (DHAP) having the ^13^C label as well at 1-C. GA3P and DHAP are combined to Fructose-1,6P (FBP) and subsequently Fructose-6P (F6P) labelled at 3-C and 4-C. During CBB cycle (**Figure 5 A**), F6P, DHAP and GA3P are combined through transaldolase and transketolase reactions, leading in the end to ribulose-5P (Ru5P) being labeled at 1-C and to similar proportions at 2-C and 3-C. RuBP is generated by PRK based on Ru5P. If RUBISCO uses 1,2,3-^13^C labeled RuBP, two molecules of 3PGA are generated, one labeled only at 1-C, and one labeled fully at 1,2,3-C. Another possibility would be the usage of gluconeogenetic reactions and upper OPP pathway (**Figure 5 B**) which should be both active during the night in *Synechocystis*. Then, F6P labeled at 3-C and 4-C is converted to gluconate-6P, which is labeled in the same pattern. Decarboxylation by gluconate-6P decarboxylase (GND) leads to Ru5P and subsequently RuBP being ^13^C labeled at 2-C and 3-C. RUBISCO carboxylase activity would then result in 3PGA labeled at 1-C and 2-C.

Based on the labeling pattern in aspartate, the regeneration via gluconeogenesis and upper OPP seems more likely. However, regeneration of one molecule RuBP costs three molecules ATP, and avoids net C_i_ assimilation due to decarboxylation by GND.

## Discussion

RUBISCO is the main carbon fixing enzyme on earth and of most photoautotrophic organisms. RUBISCO assimilates CO_2_ as part of the CBB cycle which is fueled by reduction and energy equivalents that are produced during photosynthetic light reactions. RUBISCO is regulated on several levels in vascular plants and algae, to avoid activity during the night when the energy and redox state are not favorable for such a consuming reaction. In plants, RUBISCO is inactivated by sugar inhibitors (reviewed by Orr et al. (2023)). RUBISCO activase, which is under control of thioredoxins, removes the sugar inhibitors under ATP hydrolysis (Portis et al., 1986). However, *Synechocystis* RUBISCO is neither regulated by sugar inhibitors, nor was a RUBISCO activase found in *Synechocystis* (Pearce, 2006; Lechno-Yossef et al., 2020).

In WT, there was no assimilation of CO_2_ by RUBISCO during the night (**Figure 1**), meaning that even though the mechanisms controlling RUBISCO in plants and other algae are not existing in *Synechocystis*, RUBISCO activity is successfully prevented. However, if the regulatory protein CP12, which shuts down the CBB cycle under oxidizing conditions that are prevailing during the night, is deleted, RUBISCO activity first in form of oxygenase reaction and later in from of carboxylase activity, identified by ^13^C labelling in 1-C of 3PGA, can be detected (**Figure 1**, **Figure 3**). Label in 1-C of 3PGA could hypothetically result from assimilation by PEPC in 4-C of oxaloacetate (OAA) that is interconverted in reversible reactions with malate and fumarate. As fumarate is a symmetric molecule, ^13^C would be equilibrated from 4-C to 1-C position of malate, oxaloacetate and aspartate. During gluconeogenesis, 1-C labeled oxaloacetate would be converted via PEP carboxykinase to PEP labeled at 1-C, which would be further converted by enolase and phosphoglycerate mutase to 3PGA labeled at 1-C. However, we do not see any positional swapping in WT and *Δcp12::cp12* for aspartate 1-C and 4-C even though PEPC is active (**Figure 5**, Wittemeier et al., 2024). The main flux from OAA during the night in *Synechocystis* was predicted towards aspartate (Knoop et al., 2013). Furthermore, previous studies found that fumarase in *Synechocystis* acts preferentially unidirectional towards fumarate (Hasunuma et al., 2018; Katayama et al., 2019). Consequently, we assume that label in 1-C of 3PGA is due to RUBISCO activity during the night in *Δcp12*.

RUBISCO carboxylase activity depends on two substrates: RuBP and CO_2_. In *Δcp12*, RuBP accumulates directly after onset of the night (**Figure 4**, **Figure S 6**), which can be explained by activation of glycogen catabolism via OPP pathway (Pelroy et al., 1972; Yang et al., 2002). In *Synechococcus elongatus* PCC 7942, 3PGA was labeled based on NaH^13^CO_3_ under heterotrophic conditions in darkness with glucose supply due to RuBP accumulation, when *cp12* was deleted, and even stronger when PRK and RUBISCO were overexpressed (Kanno et al., 2017). This indicates that RUBISCO activity during darkness can indeed be fueled through sugar catabolism, either of stored glycogen or externally fed glucose. In WT, we do not see high RuBP accumulation. Taken together, our assumption is that RuBP synthesis in the dark is balanced or regulated directly or indirectly through CP12 mediated inactivation of PRK (and GAPDH2). Nevertheless, we do not detect CO_2_ assimilation in *Δcp12::cp12ΔCysN* (**Figure 3**), which has a deleted cysteine binding domain for PRK. But previous studies showed that the binding to the AWD_VEEL motif also has a main role on PRK inactivation (Blanc-Garin et al., 2022). Moreover, in contrast to GAPDH2, the cyanobacterial PRK contains intrinsic cysteine pairs that allow direct redox regulation of this enzyme independently from CP12 (Fukui et al., 2022). In *Δcp12::cp12ΔCysC*, RuBP accumulated as well and CO_2_ was assimilated to a smaller amount (**Figure 4**). The complex of CP12, GAPDH2 and PRK is built sequentially. CP12 first binds GAPDH2 and subsequently PRK (McFarlane et al., 2019; Lucius et al., 2022). Potentially, PRK was not completely inactivated in this mutant, since GAPDH2 binding to CP12 might be necessary for stable PRK binding leading to free PRK molecules in the cell. In *Δcp12::cp12ΔCysNC*, CO_2_ assimilation was higher as if only the C-terminal cysteines were deleted (**Figure 3**), indicating that PRK binds less stable if the N-terminal cysteines are deleted as well. However, highest CO_2_ assimilation was found, when CP12 was deleted completely (**Figure 3**), suggesting that there is partial inactivation of GAPDH2 and PRK even without cysteine pairs. Likely, the cysteine disulfide bonds stabilize the complex but binding of GAPDH2 and PRK is partially also possible when the cysteine pairs are missing. Additionally, PRK itself is also directly redox regulated (Fukui et al., 2022).

Accumulation of RuBP directly after onset of the night correlated with ^13^C concentration of 3PGA in a time-shifted manner (**Figure 4**). RuBP concentrations decrease while 3PGA accumulates both at about 30 min (**Figure S 6)**. As RuBP is available and 2PG produced to low amounts directly after transition to darkness, RUBISCO appears to be active but may possibly lack CO_2_ in the initial phases of the pulse. Removal of oxygen from the medium did not prevent 2PG accumulation and the decrease of 3PGA (**Figure S 5**), indicating that the intracellular CO_2_ level was decreased or unavailable. The CO_2_ pulse reaches RUBISCO late and later than O_2_, which should still be present as side product from photosynthesis. The technical aspect is that under the 5% high CO_2_ conditions, we had to remove all inorganic carbon from the medium prior to the pulse to be able to monitor a rapid ^13^CO_2_ uptake. Since this was done at the end of the light period, all dissolved cellular CO_2_ was likely used up. Physiologically, we have the HCO_3_^−^ and CO_2_ uptake mechanisms down regulated under high CO_2_ conditions (Burnap et al., 2015; Spät et al., 2021). The CCM is also inactivated upon light to dark transition (Oren et al., 2021). We have evidence from our PEPC monitoring that ^13^CO_2_ arrives indeed rather late too, under our conditions, but slightly earlier than in 3PGA (**Figure 5**). This sequence is likely explained by the equilibrium of HCO_3_^−^ and CO_2_ towards accumulation of HCO_3_^−^ due to pH. Cellular pH was found to be 6.9 to 7.7 in darkness and medium with pH 8.0 (Lawrence et al., 1997; Jiang et al., 2013). Based on Henderson-Hasselbach equations, HCO_3_^−^ would be present 6 to 40-times more than CO_2_ in this pH range. Even though K_m_(CO_2_) values are slightly higher for RUBISCO (K_m_(CO_2_) ∼180-270 µM (Marcus et al., 2005; Marcus et al., 2011)) than for PEPC (K_m_(HCO_3_^−^) ∼ 800 µM (Takeya et al., 2017)), carbonic anhydrase is needed to convert HCO_3_^−^ to CO_2_ in the direct vicinity of RUBISCO as part of the CCM. Additionally, CO_2_ is not only needed as a substrate but also regulates the active state of RUBISCO by carbamylation, which has been directly verified at the *Synechocystis* RUBISCO (King et al., 2022). The highly significant correlation between RuBP concentration at 15 min and 3PGA ^13^C concentration at 90 min endorse that RUBISCO is regulated by substrate availability, in this case by RuBP availability as soon as CO_2_ is sufficiently available. This view is encouraged by the fact that RUBISCO *in vitro* activity, when RuBP and CO_2_ are unlimited, was not changed between conditions (**Figure 2**). The same applies to unchanged RUBISCO (subunit) abundance (**Figure 2**). At the end of the night, it was still possible to detect ^13^C in 1-C of 3PGA but not in 2,3-C (**Figure S 3**). It seems that at the end of the night, RuBP was still produced based on remaining glycogen catabolism. However, assimilation was much slower indicating that C_i_ uptake mechanisms were even less active at the end of the night.

In *Δcp12*, ^13^C enrichment was not only found in 1-C, but also in 2,3-C of 3PGA (**Figure 1**, **Figure 3**). This indicates that not only RUBISCO is active but also that regeneration of RuBP from 3PGA was possible (Rajarathinam et al., 2025). Regeneration via CBB cycle is a very ATP and NADPH consuming process. Since ATP and NADPH are rare during the night, because photosynthesis is shut off, RuBP regeneration via CBB cycle seems unlikely. We checked the labeling pattern of aspartate and found that 2-C but not 3-C of aspartate got labeled in *Δcp12*. 2-C and 3-C of aspartate represent 2-C and 3-C of 3PGA (Wittemeier et al., 2024) and thus, give indication of RuBP regeneration. Sole label in 2-C but not 3-C suggested regeneration of RuBP via gluconeogenesis and upper part of the OPP pathway (**Figure 5**). In general, gluconeogenetic enzyme reactions are mostly catalyzed by the same enzymes catalyzing glycolytic reactions and, hence, should be active. The OPP pathway is the main route used for sugar catabolism and also active during the night (Yang et al., 2002). This regeneration route does not consume NADPH but needs ATP as well. ATP could be provided through glycolysis and OPP pathway through phosphoglycerate kinase. However, regeneration via gluconeogenesis and OPP does not only consume ATP but also does not lead to net CO_2_ assimilation, since one CO_2_ molecule per assimilated CO_2_ is lost via GND.

In conclusion, this study revealed that RUBISCO activity during the night is regulated by substrate provision in *Synechocystis*. Application of ^13^C positional analysis not only uncovered that RUBISCO assimilates CO_2_ during the night in *Δcp12*, but also that regeneration of RuBP is possible likely by using gluconeogenetic and OPP pathways. This means that RUBISCO itself is not per se inactivated, but both substrates, CO_2_ and RuBP, are heavily reduced through downregulation of CCM and inactivation of regeneration enzymes, respectively. The functions of the Cys binding domains were discussed and importance of both in allowing stable complex formation with GAPDH2 and PRK was demonstrated. A CP12 version without Cys binding domains did not fully constitute the deletion of *cp12*, suggesting that the protein backbone also has regulatory functions on GAPDH2 and PRK binding.

## Materials and methods

### Cultivation and sampling of *Synechocystis* sp. PCC 6803

All experiments were conducted with the glucose-tolerant wild type (WT) of *Synechocystis* sp. PCC 6803 (*Synechocystis*) and respective *cp12* deletion mutant (*Δcp12*) and complementation mutants (*Δcp12::cp12*) generated by Lucius et al. (2022). All mutant strains were grown by adding the respective antibiotics (*Δcp12*: 50 µg/mL kanamycin, *Δcp12::cp12*: 50 µg/mL kanamycin and 40 µg/mL spectinomycin).

Cultivations and ^13^CO_2_ labeling experiments were conducted as described before (Wittemeier et al., 2024). Briefly*, Synechocystis* WT and mutants were cultivated photoautotrophically in modified BG11 growth medium (Rippka et al., 1979) without citric acid and with FeCl_3_ instead of ferric citrate at 30°C. Cultures were constantly bubbled with 5% CO_2_-enriched air. Cells were grown for 3 days in constant light (100 µmol photons m^−2^ s^−1^). Medium was renewed by centrifugation and illumination was changed to 12 h light/12 h dark cycles (100 µmol photons m^−2^ s^−1^ in light phases) and cultivated for 3 more days. Medium was renewed 4 h prior to the fourth night by fast vacuum filtration and continuous illumination and OD adjusted to ∼0.8. Cells were sampled by <15 s filtration onto glass fiber filters (25 mm, pore size 1.2 µm, Cytiva, Sigma Aldrich/Merck KGaA) and immediate shock freezing in liquid N_2_. Directly before transition to the fourth night, t_0_ was harvested in the light with 5% ambient CO_2_ bubbling. Culture medium was exchanged by fast filtration to remove dissolved non-labeled Ci from the cultures. Cells were resuspended in the dark by immediate bubbling with 5% ^13^CO_2_ in synthetic air. Labeled samples at t_1_-t_6_ were collected 5, 10, 15, 30, 60 and 90 min after beginning of ^13^CO_2_ bubbling. OD_750_ mL^−1^ was adjusted to ∼1.0 and recorded of all individual cultures at t_0_, t_1_ and t_6_. For labeling studies at the end of the dark period and at transition from darkness to light (night 4 to day 5), procedure was the same except having all timepoints and media exchanges in complete darkness or starting 5% ^13^CO_2_ labeling after onset of the day, respectively.

### Metabolite extraction, derivatization and quantification by GC-MS

Extraction of polar metabolites from deep-frozen cells on filters was done using methanol (Sigma-Aldrich, gradient grade for liquid chromatography, ≥99.9%), chloroform (Sigma-Aldrich/Merck KGaA, contains ethanol as stabilizer, ACS reagent grade, ≥99.8%) and double distilled water (ddH_2_O) as described before (Wittemeier et al., 2024). In short, filters were incubated with 1 mL of extraction mix (methanol:chloroform:ddH_2_O; 2.5:1:1; v/v/v) with 6 µg * mL^−1 13^C_6_-sorbitol as internal standard at 70°C. Phase separation was induced by adding 400 µL of ddH_2_O. The polar phase was dried overnight by vacuum centrifugation and used for GC-MS analysis.

Chemical derivatization of dried metabolite samples for GC-MS analysis was exactly as described previously omitting 4-(dimethylamino)pyridine (Erban et al., 2020). Samples were subjected to methoxyamination followed by trimethylsilylation (TMS) using *N,O*-bis(trimethylsilyl)trifluoroacetamide (BSTFA, Macherey-Nagel, Düren, Germany).

Derivatized samples were analyzed by an Agilent 6890N24 gas chromatograph (Agilent Technologies, Waldbronn, Germany) hyphenated to either electron impact ionization-time of flight-mass spectrometry (EI-TOF-MS) using a LECO Pegasus III time of flight mass spectrometer (LECO Instrumente GmbH, Mönchengladbach, Germany) or to atmospheric pressure chemical ionization-time of flight-mass spectrometry (APCI-TOF-MS) with a micrOTOF-Q II hybrid quadrupole time-of-flight mass spectrometer (Bruker Daltonics, Bremen, Germany) equipped with an APCI ion source and GC interface (Bruker Daltonics) (Kopka et al., 2017). All measurements were conducted in splitless mode using 5% phenyl - 95% dimethylpolysiloxane fused silica capillary column with 30 m length, 0.25 mm inner diameter, 0.25 µm film thickness and an integrated 10 m precolumn (Agilent Technologies (CP9013)). Retention index standardization was based on *n*-alkanes as described earlier (Erban et al., 2020).

GC-EI-MS chromatograms were recorded at nominal mass resolution, baseline corrected and processed as described previously (Erban et al., 2020). Chemical reference compounds and their analytes were picked by manual supervision using TagFinder (Luedemann et al., 2011) and the NIST MS Search 2.0 software (http://chemdata.nist.gov/). Observed experimental mass spectra and retention time indices (RI) were matched to the mass spectral and RI reference collection of the Golm Metabolome Database (GMD; Kopka et al., 2005). A######-### identifiers directly relate to GMD entries. Quantification of isotopologues and isotopologue distributions (MIDs) were based on peak apex abundances.

Exact mass of GC-APCI-MS files were internally calibrated based on PFTBA (Kopka et al., 2017). Files were transcribed into mzXML format using Bruker DataAnalysis and AutomationEngine software (version 4.2). Analytes of GC-APCI-MS files were identified manually based on exact monoisotopic masses, comparison to the paired GC-EI-MS analyses and parallel measurements of metabolite reference compounds. The isotopologue abundances of molecular ions and mass fragments and respective ^13^C labeled MIDs were extracted from each GC-APCI-MS files in a defined chromatographic time range adjusted to each analyte and in a mass range of ± 0.005 mass units using the R packages xcms (version 3.22.0; Tautenhahn et al., 2008), MSnbase (version 2.26.0; Gatto et al., 2021) and msdata (version 0.40.0; Neumann and Gatto, 2017) in RStudio (2023.6.1.524, http://www.posit.co/, R version 4.3.1). Quantification of isotopologue abundances was based on peak area under the peak apex ± 10 scans.

For metabolite quantification, the sum of all isotopologue abundances was used. Metabolite concentration was normalized to internal standard ^13^C_6_-Sorbitol, OD_750_ and sample volume. Molar metabolite concentrations were acquired through parallel analysis of calibration series of non-labeled reference compounds.

### Metabolite extraction and quantification by LC-MS/MS

Sugar phosphates and triose phosphates were extracted from frozen cells on glass fiber filters by cold methanol chloroform water extraction. Briefly, 1 mL 90% (v/v) methanol (−20°C) and 260 µL chloroform were added to the frozen cells on liquid N_2_. Cells were broken by 3 freeze-thaw cycles. Phase separation was induced by adding 600 µL ddH_2_O and the polar phase collected for 3 times. Extracts were freeze-dried over night by lyophilization. Dried extracts were resuspended in ddH_2_O and filtered for subsequent LC-MS/MS analysis. LC-MS/MS quantification was performed as described in Medeiros et al. (2022) using ¼ diluted and undiluted extracts.

### Calculation of E^13^C and positional E^13^C analysis

MIDs and resulting ^13^C enrichment calculations of mass features were corrected for natural isotope abundances (NIA) according to their specific molecular formula using RStudio and the IsoCorrectoR package (version 1.18.0; Heinrich et al., 2018). Correction of tracer impurity was done manually by adjusting IsoCorrectoR results to the specified ^13^C purity of each reference substance. E^13^C and molar concentrations of 3PGA and aspartic acid were used to calculate molar ^13^C concentration at each carbon position of the 3PGA and aspartic acid backbone (Wittemeier et al., 2024; Rajarathinam et al., 2025).

### Western blot analysis

Cells corresponding to OD_750_=4 were resuspended in 400 µL PBS buffer (140 mM NaCl, 10 mM phosphate buffer, 3mM KCl, pH 7.4) and lysed by agitation with glass beads at 30 Hz for 10 min at 4°C followed by centrifugation for 2 min at 2,000 xg and 4°C. The protein content of the supernatant was determined using the Pierce™ BCA Protein Assay Kit (Thermo Fisher Scientific). Proteins were mixed with 4x Laemmli sample buffer (0.2 M TRIS-HCl pH 6.5, 0.4 M DTT, 8% SDS, 32% glycerol, bromphenolblue) and denatured for 10 min at 90°C. Separation was done by SDS-PAGE (8% stacking gel, 12% resolving; Laemmli, 1970). Gels were stained with Quick Coomassie Stain (Serva). Proteins were blotted onto a PVDF membrane by wet Western transfer. Membranes were subsequently blocked with 5% milk in TBS-T. Detection of RUBISCO was facilitated by RbcL antibody (Agisera AS03037) followed by anti-Rabbit IgG (H+L) HRP-conjugated (Invitrogen). For illumination, PIERCE ECL Western Blotting Substrate (Thermo Fisher Scientific) was used.

### Proteomics

*Synechocystis* cultures were grown to an OD_750_ of 1 and 25 mL were harvested by centrifugation shortly before and 90 min after onset of the night. The cell pellet was dissolved in 1.5 mL SDS buffer (4% SDS, 100 mM TRIS/HCl pH 8.0) and incubated at 95°C for 10 min. DNA comminution was done by sonication (Bandelin ultrasonic homogenizer HD 2070.2) for 1 min (40% pulse time, amplitude 50%). Protein disulfide bonds were reduced with 10 mM DTT for 45 min. Reduced cysteine thiol groups were alkylated with 5.5 mM IAA for 45 min. Proteins were precipitated with 9 sample volumes acetone/methanol (8:1) mixture, centrifuged and washed with 80% acetone. Protein was dissolved in 200 µL 10 mM TRIS/HCl pH 7.5 and freeze-dried overnight.

For LC-MS/MS analysis, samples were resolubilized in denaturation buffer (6 M urea, 2 M thiourea in 100 mM Tris/HCl; pH 8.0) at a final protein concentration of 1 μg/μL. Protein disulfide bonds were reduced with 1 mM DTT for 45 min and the resulting thiol groups were alkylated with iodoacetamide at a final concentration of 5.5 mM for 45 min. Predigestion of proteins with Lys-C (Santa Cruz Biotechnology) for 3 h was followed by dilution with 4 volumes of 20 mM ammonium bicarbonate buffer, pH 8, and overnight digestion with trypsin (sequencing grade, Promega), both at protease/protein ratios of 1/100. The resulting peptide solutions were acidified with trifluoroacetic acid to pH 2.5, and an aliquot corresponding to 10 μg peptides was purified by stage tips (Ishihama et al., 2006). For LC‒MS/MS-based protein analysis, 500 ng of each sample was loaded onto an in-house-made 20 cm column with 75 μm ID, packed with ReproSil-Pur 1.9 μm C18 material (Dr. Maisch, Germany) and separated by reverse phase chromatography on an EASY-nLC 1200 system (Thermo Fisher Scientific, USA) using 90 min gradients. Eluting peptides were analyzed on an Orbitrap Exploris 480 mass spectrometer (Thermo Fisher Scientific, USA) running in data dependent acquisition mode. All raw spectra were processed with MaxQuant software (version 2.4.2.0) with the following parameters: Carbamidomethyl (C) as a fixed modification; oxidation (M) and N-terminal acetylation as variable modifications; digestion mode, trypsin/P, with up to two missed cleavages. The false discovery rate (FDR) was set to 0.01 at both the peptide and protein levels. The “match between runs” option was enabled (Cox and Mann, 2008). The acquired m/z spectra were searched against the proteome databases of *Synechocystis* sp. PCC 6803 (downloaded from Uniprot). Subsequent data analysis was performed using the Perseus software suite (version 1.6.5.0; Tyanova et al., 2016).

### Determination of RUBISCO *in vitro* carboxylase activity

Total soluble protein was extracted in RUBISCO activation buffer (20 mM Bicine/NaOH, pH 8.0, 50 mM MgCl_2_, 50 mM NaHCO_3_) from 20 mL *Synechocystis* culture through vortexing with glass beads (150-300 mm). Protein quantification was done by amido black staining and absorbance measurements with the LAMBDA 365+ UV/Vis Spectrophotometer (PerkinElmer; Schulz et al., 1994). RUBISCO was activated by incubation for 1 h at room temperature. Carboxylase activity was determined using 30 µg of total protein in assay buffer (100 mM Bicine/KOH, pH 8.2, 20 mM MgCl_2_, 33 mM NaH^14^CO_3_, and 0.66 mM RuBP) at 25°C (Timm et al., 2015). Total activity was determined after activation for 5 min at 25°C in the absence of RuBP and initiated by adding RuBP. The reaction was quenched after 60, 90, 120, and 150 s by adding the same volume of 10 M formic acid. Samples were dried at 80°C to remove all inorganic labeled carbon, rehydrated with 500 µL H_2_O and mixed with 5 mL liquid scintillation cocktail (Ultima Gold; PerkinElmer). Amount of C^14^ was quantified by liquid scintillation counting (Liquid Scintillation Analyzer Tri-Carb 2810 TR; PerkinElmer).

## Acknowledgements

We acknowledge the funding of the German Research Foundation (DFG) in the framework of the research consortium SCyCode (FOR2816; KO 2329/7-2, HA 2002/23-2, MA 4918/4-2) and the support of the Max Planck Society. We thank Prof. Dr. Mark Stitt for helpful discussions.

## Competing interests

The authors declare no potential conflict of interest.

## Author contributions

J.K. and M.H. conceived the original research plan. L.W. designed and performed the experiments. L.W. performed cyanobacteria cultivation and labelling studies, Western Blot analysis and GC-MS analysis. S.A. performed the LC-MS analysis. N.S. performed RUBISCO *in vitro* assays. N.N. and B.M. did the proteome analysis. J.K. and L.W. wrote the manuscript with contribution of all other authors.

## Data availability

This study does not include large-scale datasets. The data supporting the findings of this study are presented in Figs 1-5 and Figs S1-S7 of this article.

## Notes

### Competing Interest Statement

The authors have declared no competing interest.

## References

Bar-On, Y.M., and Milo, R. (2019). The global mass and average rate of rubisco. Proc Natl Acad Sci U S A 116, 4738–4743.

Berwanger, L.C., Thumm, N., Stirba, F.P., Gholamipoorfard, R., Pawlowski, A., Kolkhof, P., Volke, J., Kollmann, M., Wiegard, A., and Axmann, I.M. (2025). Self-sustained rhythmic behavior of Synechocystis sp. PCC 6803 under continuous light conditions in the absence of light–dark entrainment. PNAS Nexus 4, pgaf120.

Blanc-Garin, V., Veaudor, T., Setif, P., Gontero, B., Lemaire, S.D., Chauvat, F., and Cassier-Chauvat, C. (2022). First in vivo analysis of the regulatory protein CP12 of the model cyanobacterium Synechocystis PCC 6803: Biotechnological implications. Front Plant Sci 13, 999672.

Burnap, R.L., Hagemann, M., and Kaplan, A. (2015). Regulation of CO2 Concentrating Mechanism in Cyanobacteria. Life (Basel) 5, 348–371.

Cleland, W.W., Andrews, T.J., Gutteridge, S., Hartman, F.C., and Lorimer, G.H. (1998). Mechanism of Rubisco: The Carbamate as General Base. Chem Rev 98, 549–562.

Cox, J., and Mann, M. (2008). MaxQuant enables high peptide identification rates, individualized p.p.b.-range mass accuracies and proteome-wide protein quantification. Nat Biotechnol 26, 1367–1372.

Dismukes, G.C., Klimov, V.V., Baranov, S.V., Kozlov, Y.N., DasGupta, J., and Tyryshkin, A. (2001). The origin of atmospheric oxygen on Earth: the innovation of oxygenic photosynthesis. Proc Natl Acad Sci U S A 98, 2170–2175.

Erban, A., Martinez-Seidel, F., Rajarathinam, Y., Dethloff, F., Orf, I., Fehrle, I., Alpers, J., Beine-Golovchuk, O., and Kopka, J. (2020). Multiplexed profiling and data processing methods to identify temperature-regulated primary metabolites using gas chromatography coupled to mass spectrometry. In Plant Cold Acclimation, D.K. Hincha and E. Zuther, eds (New York, NY: Springer US), pp. 203–239.

Flecken, M., Wang, H., Popilka, L., Hartl, F.U., Bracher, A., and Hayer-Hartl, M. (2020). Dual Functions of a Rubisco Activase in Metabolic Repair and Recruitment to Carboxysomes. Cell 183, 457–473.e420.

Fukui, K., Yoshida, K., Yokochi, Y., Sekiguchi, T., Wakabayashi, K.I., Hisabori, T., and Mihara, S. (2022). The Importance of the C-Terminal Cys Pair of Phosphoribulokinase in Phototrophs in Thioredoxin-Dependent Regulation. Plant Cell Physiol 63, 855–868.

Gatto, L., Gibb, S., and Rainer, J. (2021). MSnbase, efficient and elegant R-based processing and visualization of raw mass spectrometry data. Journal of Proteome Research 20, 1063–1069.

Groben, R., Kaloudas, D., Raines, C.A., Offmann, B., Maberly, S.C., and Gontero, B. (2010). Comparative sequence analysis of CP12, a small protein involved in the formation of a Calvin cycle complex in photosynthetic organisms. Photosynthesis Research 103, 183–194.

Gurrieri, L., Fermani, S., Zaffagnini, M., Sparla, F., and Trost, P. (2021). Calvin-Benson cycle regulation is getting complex. Trends Plant Sci 26, 898–912.

Hagemann, M., and Hess, W.R. (2018). Systems and synthetic biology for the biotechnological application of cyanobacteria. Curr Opin Biotechnol 49, 94–99.

Hagemann, M., Song, S., and Brouwer, E.-M. (2021). Inorganic Carbon Assimilation in Cyanobacteria: Mechanisms, Regulation, and Engineering. In Cyanobacteria Biotechnology, pp. 1–31.

Hasunuma, T., Matsuda, M., Kato, Y., Vavricka, C.J., and Kondo, A. (2018). Temperature enhanced succinate production concurrent with increased central metabolism turnover in the cyanobacterium *Synechocystis* sp. PCC 6803. Metabolic Engineering 48, 109–120.

Hauf, W., Watzer, B., Roos, N., Klotz, A., and Forchhammer, K. (2015). Photoautotrophic Polyhydroxybutyrate Granule Formation Is Regulated by Cyanobacterial Phasin PhaP in Synechocystis sp. Strain PCC 6803. Appl Environ Microbiol 81, 4411–4422.

Heinrich, P., Kohler, C., Ellmann, L., Kuerner, P., Spang, R., Oefner, P.J., and Dettmer, K. (2018). Correcting for natural isotope abundance and tracer impurity in MS-, MS/MS- and high-resolution-multiple-tracer-data from stable isotope labeling experiments with IsoCorrectoR. Scientific Reports 8, 17910.

Ishihama, Y., Rappsilber, J., and Mann, M. (2006). Modular stop and go extraction tips with stacked disks for parallel and multidimensional Peptide fractionation in proteomics. J Proteome Res 5, 988–994.

Jiang, H.B., Cheng, H.M., Gao, K.S., and Qiu, B.S. (2013). Inactivation of Ca(2+)/H(+) exchanger in Synechocystis sp. strain PCC 6803 promotes cyanobacterial calcification by upregulating CO(2)-concentrating mechanisms. Appl Environ Microbiol 79, 4048–4055.

Kanno, M., Carroll, A.L., and Atsumi, S. (2017). Global metabolic rewiring for improved CO(2) fixation and chemical production in cyanobacteria. Nat Commun 8, 14724.

Katayama, N., Takeya, M., and Osanai, T. (2019). Biochemical characterisation of fumarase C from a unicellular cyanobacterium demonstrating its substrate affinity, altered by an amino acid substitution. Scientific Reports 9, 10629.

King, D.T., Zhu, S., Hardie, D.B., Serrano-Negrón, J.E., Madden, Z., Kolappan, S., and Vocadlo, D.J. (2022). Chemoproteomic identification of CO2-dependent lysine carboxylation in proteins. Nature Chemical Biology 18, 782–791.

Knoop, H., Grundel, M., Zilliges, Y., Lehmann, R., Hoffmann, S., Lockau, W., and Steuer, R. (2013). Flux balance analysis of cyanobacterial metabolism: the metabolic network of *Synechocystis* sp. PCC 6803. PLOS Computational Biology 9, e1003081.

Köbler, C., Schmelling, N.M., Wiegard, A., Pawlowski, A., Pattanayak, G.K., Spat, P., Scheurer, N.M., Sebastian, K.N., Stirba, F.P., Berwanger, L.C., Kolkhof, P., Macek, B., Rust, M.J., Axmann, I.M., and Wilde, A. (2024). Two KaiABC systems control circadian oscillations in one cyanobacterium. Nat Commun 15, 7674.

Koch, M., Berendzen, K.W., and Forchhammer, K. (2020). On the Role and Production of Polyhydroxybutyrate (PHB) in the Cyanobacterium Synechocystis sp. PCC 6803. In Life.

Koksharova, O., Schubert, M., Shestakov, S., and Cerff, R. (1998). Genetic and biochemical evidence for distinct key functions of two highly divergent GAPDH genes in catabolic and anabolic carbon flow of the cyanobacterium Synechocystis sp. PCC 6803. Plant Mol Biol 36, 183–194.

Kopka, J., Schauer, N., Krueger, S., Birkemeyer, C., Usadel, B., Bergmuller, E., Dormann, P., Weckwerth, W., Gibon, Y., Stitt, M., Willmitzer, L., Fernie, A.R., and Steinhauser, D. (2005).: the Golm Metabolome Database. Bioinformatics 21, 1635–1638.

Kopka, J., Schmidt, S., Dethloff, F., Pade, N., Berendt, S., Schottkowski, M., Martin, N., Duhring, U., Kuchmina, E., Enke, H., Kramer, D., Wilde, A., Hagemann, M., and Friedrich, A. (2017). Systems analysis of ethanol production in the genetically engineered cyanobacterium *Synechococcus* sp. PCC 7002. Biotechnology for Biofuels and Bioproducts 10, 56.

Laemmli, U.K. (1970). Cleavage of structural proteins during the assembly of the head of bacteriophage T4. Nature 227, 680–685.

Lawrence, B.A., Polse, J., DePina, A., Allen, M.M., and Kolodny, N.H. (1997). 31P NMR identification of metabolites and pH determination in the cyanobacterium Synechocystis sp. PCC 6308. Curr Microbiol 34, 280–283.

Lechno-Yossef, S., Rohnke, B.A., Belza, A.C.O., Melnicki, M.R., Montgomery, B.L., and Kerfeld, C.A. (2020). Cyanobacterial carboxysomes contain an unique rubisco-activase-like protein. New Phytologist 225, 793–806.

Lopez-Calcagno, P.E., Howard, T.P., and Raines, C.A. (2014). The CP12 protein family: a thioredoxin-mediated metabolic switch? Frontiers in Plant Science Volume 5–2014.

López-Calcagno, P.E., Abuzaid, A.O., Lawson, T., and Raines, C.A. (2017). Arabidopsis CP12 mutants have reduced levels of phosphoribulokinase and impaired function of the Calvin–Benson cycle. Journal of Experimental Botany 68, 2285–2298.

Lorimer, G.H., Badger, M.R., and Andrews, T.J. (1976). The activation of ribulose-1,5-bisphosphate carboxylase by carbon dioxide and magnesium ions. Equilibria, kinetics, a suggested mechanism, and physiological implications. Biochemistry 15, 529–536.

Lucius, S., and Hagemann, M. (2024). The primary carbon metabolism in cyanobacteria and its regulation. Front Plant Sci 15, 1417680.

Lucius, S., Theune, M., Arrivault, S., Hildebrandt, S., Mullineaux, C.W., Gutekunst, K., and Hagemann, M. (2022). CP12 fine-tunes the Calvin-Benson cycle and carbohydrate metabolism in cyanobacteria. Front Plant Sci 13, 1028794.

Luedemann, A., von Malotky, L., Erban, A., and Kopka, J. (2011). TagFinder: preprocessing software for the fingerprinting and the profiling of gas chromatography–mass spectrometry based metabolome analyses. In Plant Metabolomics, N.W. Hardy and R.D. Hall, eds (Totowa, NJ: Humana Press), pp. 255–286.

Makowka, A., Nichelmann, L., Schulze, D., Spengler, K., Wittmann, C., Forchhammer, K., and Gutekunst, K. (2020). Glycolytic shunts replenish the Calvin-Benson-Bassham cycle as anaplerotic reactions in cyanobacteria. Molecular Plant 13, 471–482.

Marcus, Y., and Gurevitz, M. (2000). Activation of cyanobacterial RuBP-carboxylase/oxygenase is facilitated by inorganic phosphate via two independent mechanisms. Eur J Biochem 267, 5995–6003.

Marcus, Y., Altman-Gueta, H., Finkler, A., and Gurevitz, M. (2005). Mutagenesis at two distinct phosphate-binding sites unravels their differential roles in regulation of Rubisco activation and catalysis. J Bacteriol 187, 4222–4228.

Marcus, Y., Altman-Gueta, H., Wolff, Y., and Gurevitz, M. (2011). Rubisco mutagenesis provides new insight into limitations on photosynthesis and growth in Synechocystis PCC6803. Journal of Experimental Botany 62, 4173–4182.

Margulis, L. (1970). Origin of Eukaryotic Cells: Evidence and Research Implications for a Theory of the Origin and Evolution of Microbial, Plant, and Animal Cells on the Precambrian Earth. (Yale University Press).

Marri, L., Thieulin-Pardo, G., Lebrun, R., Puppo, R., Zaffagnini, M., Trost, P., Gontero, B., and Sparla, F. (2014). CP12-mediated protection of Calvin–Benson cycle enzymes from oxidative stress. Biochimie 97, 228–237.

McFarlane, C.R., Shah, N.R., Kabasakal, B.V., Echeverria, B., Cotton, C.A.R., Bubeck, D., and Murray, J.W. (2019). Structural basis of light-induced redox regulation in the Calvin-Benson cycle in cyanobacteria. Proc Natl Acad Sci U S A 116, 20984–20990.

Medeiros, D.B., Ishihara, H., Guenther, M., Rosado de Souza, L., Fernie, A.R., Stitt, M., and Arrivault, S. (2022). 13CO2 labeling kinetics in maize reveal impaired efficiency of C4 photosynthesis under low irradiance. Plant Physiology 190, 280–304.

Neumann, S., and Gatto, L. (2017). msdata: Various Mass Spectrometry raw data example files., pp. R package.

Oren, N., Timm, S., Frank, M., Mantovani, O., Murik, O., and Hagemann, M. (2021). Red/far-red light signals regulate the activity of the carbon-concentrating mechanism in cyanobacteria. Sci Adv 7.

Orr, D.J., Robijns, A.K.J., Baker, C.R., Niyogi, K.K., and Carmo-Silva, E. (2023). Dynamics of Rubisco regulation by sugar phosphate derivatives and their phosphatases. J Exp Bot 74, 581–590.

Pearce, F.G. (2006). Catalytic by-product formation and ligand binding by ribulose bisphosphate carboxylases from different phylogenies. Biochemical Journal 399, 525–534.

Pelroy, R.A., Rippka, R., and Stanier, R.Y. (1972). Metabolism of glucose by unicellular blue-green algae. Arch Mikrobiol 87, 303–322.

Portis, A.R., Salvucci, M.E., and Ogren, W.L. (1986). Activation of Ribulosebisphosphate Carboxylase/Oxygenase at Physiological CO(2) and Ribulosebisphosphate Concentrations by Rubisco Activase. Plant Physiol 82, 967–971.

Price, G.D., Badger, M.R., Woodger, F.J., and Long, B.M. (2008). Advances in understanding the cyanobacterial CO_2_-concentrating-mechanism (CCM): functional components, Ci transporters, diversity, genetic regulation and prospects for engineering into plants. Journal of Experimental Botany 59, 1441–1461.

Rajarathinam, Y., Wittemeier, L., Gutekunst, K., Hagemann, M., and Kopka, J. (2025). Dynamic photosynthetic labeling and carbon-positional mass spectrometry monitor *in vivo* RUBISCO carbon assimilation rates. Plant Physiology 197.

Raven, J.A., Cockell, C.S., and De La Rocha, C.L. (2008). The evolution of inorganic carbon concentrating mechanisms in photosynthesis. Philos Trans R Soc Lond B Biol Sci 363, 2641–2650.

Rippka, R., Deruelles, J., Waterbury, J.B., Herdman, M., and Stanier, R.Y. (1979). Generic assignments, strain histories and properties of pure cultures of cyanobacteria. Microbiology 111, 1–61.

Schulz, J., Dettlaff, S., Fritzsche, U., Harms, U., Schiebel, H., Derer, W., Fusenig, N.E., Hulsen, A., and Bohm, M. (1994). The amido black assay: a simple and quantitative multipurpose test of adhesion, proliferation, and cytotoxicity in microplate cultures of keratinocytes (HaCaT) and other cell types growing adherently or in suspension. J Immunol Methods 167, 1–13.

Spät, P., Barske, T., Maček, B., and Hagemann, M. (2021). Alterations in the CO_2_ availability induce alterations in the phosphoproteome of the cyanobacterium Synechocystis sp. PCC 6803. New Phytol. 231, 1123–1137.

Stanley, D.N., Raines, C.A., and Kerfeld, C.A. (2013). Comparative Analysis of 126 Cyanobacterial Genomes Reveals Evidence of Functional Diversity Among Homologs of the Redox-Regulated CP12 Protein Plant Physiology 161, 824–835.

Takeya, M., Hirai, M.Y., and Osanai, T. (2017). Allosteric Inhibition of Phosphoenolpyruvate Carboxylases is Determined by a Single Amino Acid Residue in Cyanobacteria. Sci Rep 7, 41080.

Tamoi, M., Miyazaki, T., Fukamizo, T., and Shigeoka, S. (2005). The Calvin cycle in cyanobacteria is regulated by CP12 via the NAD(H)/NADP(H) ratio under light/dark conditions. The Plant Journal 42, 504–513.

Tautenhahn, R., Bottcher, C., and Neumann, S. (2008). Highly sensitive feature detection for high resolution LC/MS. BMC Bioinformatics 9, 504.

Timm, S., Wittmiss, M., Gamlien, S., Ewald, R., Florian, A., Frank, M., Wirtz, M., Hell, R., Fernie, A.R., and Bauwe, H. (2015). Mitochondrial dihydrolipoyl dehydrogenase activity shapes photosynthesis and photorespiration of *Arabidopsis thaliana*. Plant Cell 27, 1968–1984.

Tyanova, S., Temu, T., Sinitcyn, P., Carlson, A., Hein, M.Y., Geiger, T., Mann, M., and Cox, J. (2016). The Perseus computational platform for comprehensive analysis of (prote)omics data. Nat Methods 13, 731–740.

Wedel, N., and Soll, J. (1998). Evolutionary conserved light regulation of Calvin cycle activity by NADPH-mediated reversible phosphoribulokinase/CP12/ glyceraldehyde-3-phosphate dehydrogenase complex dissociation. Proceedings of the National Academy of Sciences 95, 9699–9704.

Wittemeier, L., Rajarathinam, Y., Erban, A., Hagemann, M., and Kopka, J. (2024). Positional ^13^C Enrichment Analysis of Aspartate by GC-MS to Determine PEPC Activity *In Vivo*. bioRxiv.

Yang, C., Hua, Q., and Shimizu, K. (2002). Metabolic flux analysis in *Synechocystis* using isotope distribution from ^13^C-labeled glucose. Metabolic Engineering 4, 202–216.

