## Supplemental Figures S1-S7 for "*In vivo* RUBISCO activity in *Synechocystis* is regulated by RuBP availability"

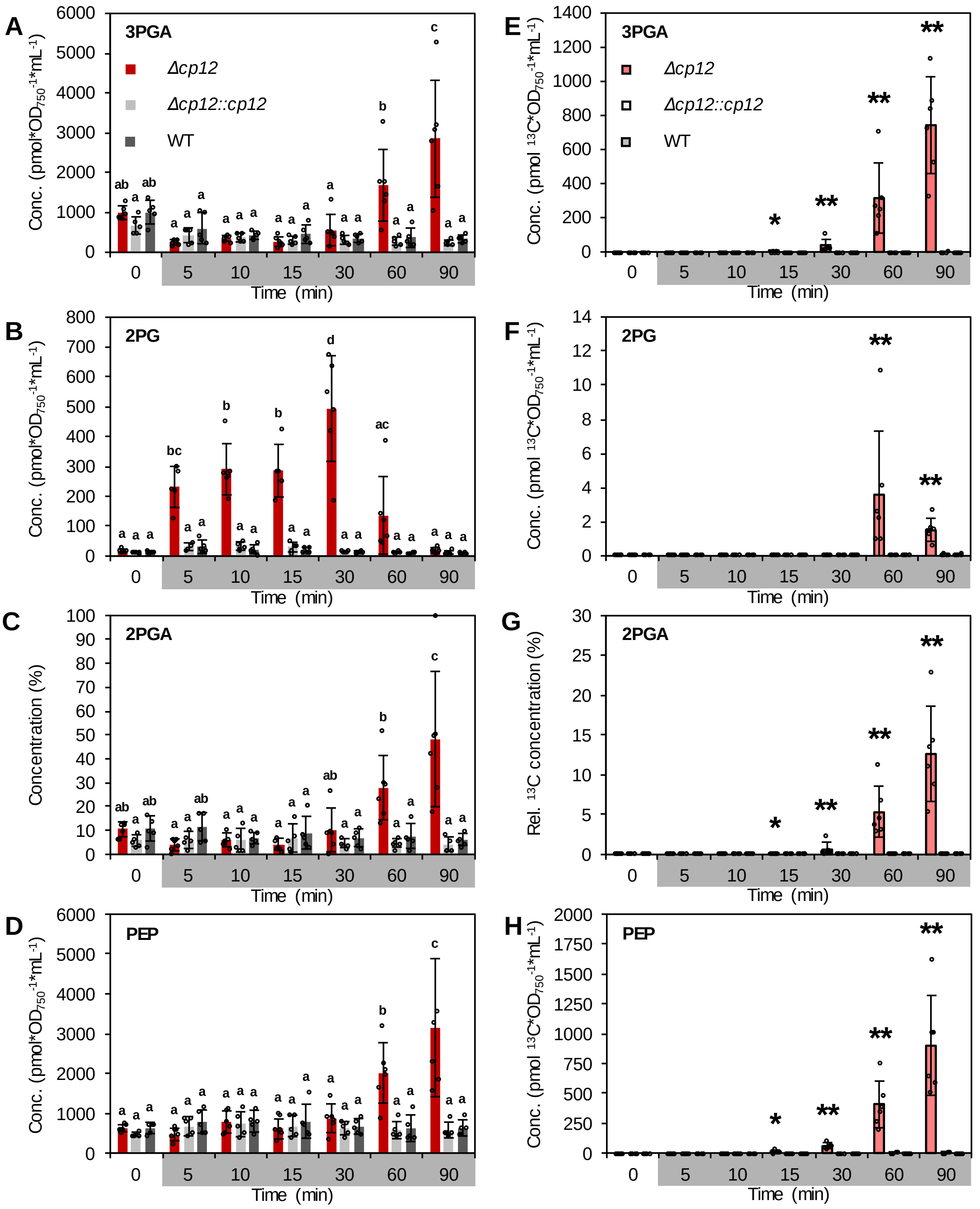

**Figure S 1. E$^{13}$C and metabolite concentration analysis of 3PGA, 2PG, 2PGA and PEP in *Δcp12* compared to WT and *Δcp12::cp12*.** *Synechocystis Δcp12* was cultivated in comparison to *Δcp12::cp12* and WT. Sampling and $^{13}CO_2$ labeling was done at transition to the fourth night. Metabolite levels (**A-D**) were determined using GC-EI-MS, $^{13}$C enrichment using GC-APCI-MS (**E-H**). 3PGA (**A, E**), 2PG (**B, F**) and PEP (**D, H**) were quantified absolutely by standard measurements. Relative concentration analysis (maximum-scaled relative intensity) was done for 2PGA (**C, G**). Bars show the average of 5 independent biological replicates ± standard deviation. Asterisks show significant differences between *Δcp12* and WT and *Δcp12::cp12* (tested independently) based on Wilcoxon-Mann-Whitney testing (*: $p<0.05$, **: $p<0.01$). Letters display significant differences of 3PGA concentration between all conditions based on Tukey's HSD ($p<0.05$).

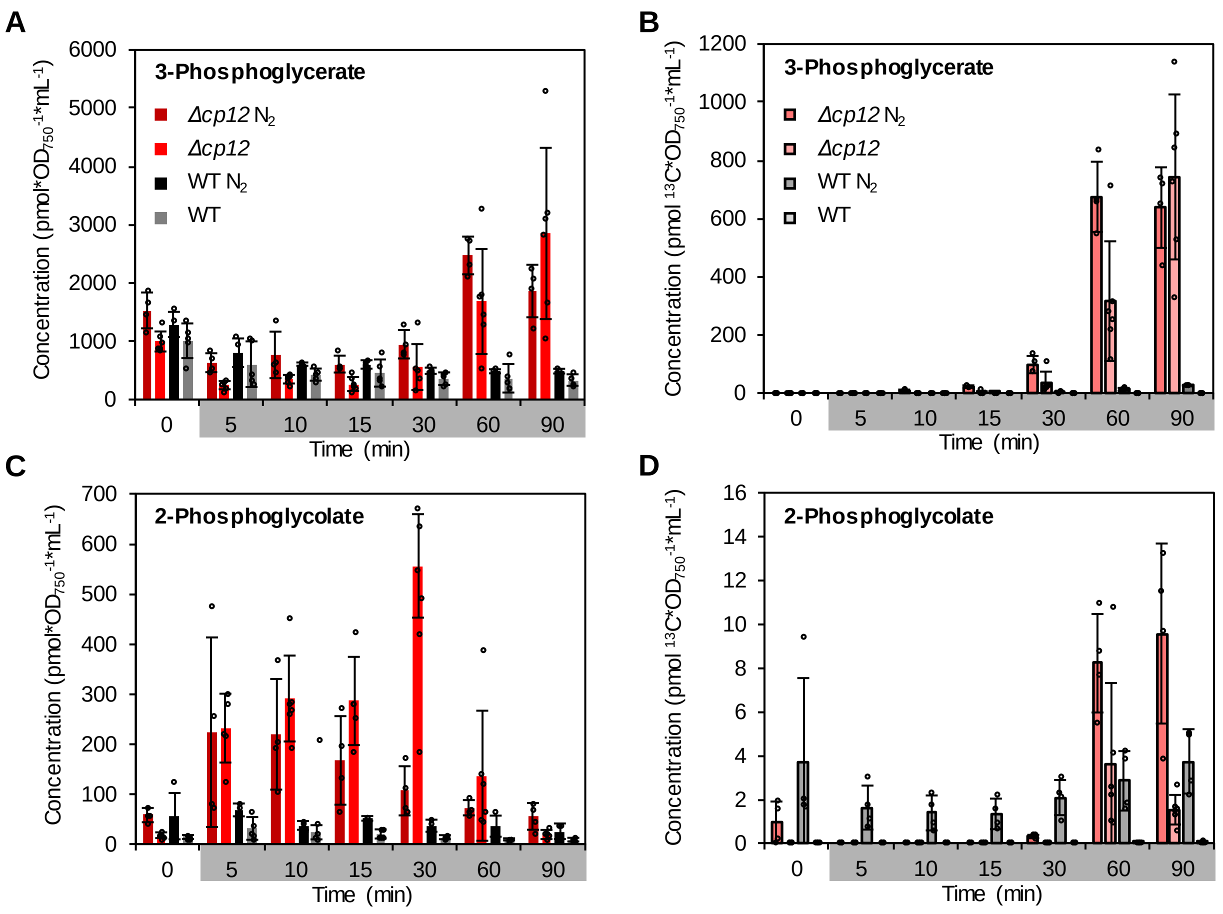

**Figure S 2. Removal of dissolved oxygen in the medium does not influence accumulation and $^{13}C$-enrichment of RUBISCO products.** Comparison of 3PGA (**A+B**) and 2PG (**C+D**) metabolite abundances in WT and *Δcp12* transferred to either normal medium or $N_2$ pre-saturated medium ($N_2$) to verify if the oxygen concentration in the medium has an influence. Metabolite concentration (pmol*$OD_{750}$*$mL^{-1}$) of 3PGA (**A**) and 2PG (**C**) and $E^{13}C$ concentration (pmol*$OD_{750}$*$mL^{-1}$) in 3PGA (**B**) and 2PG (**D**).

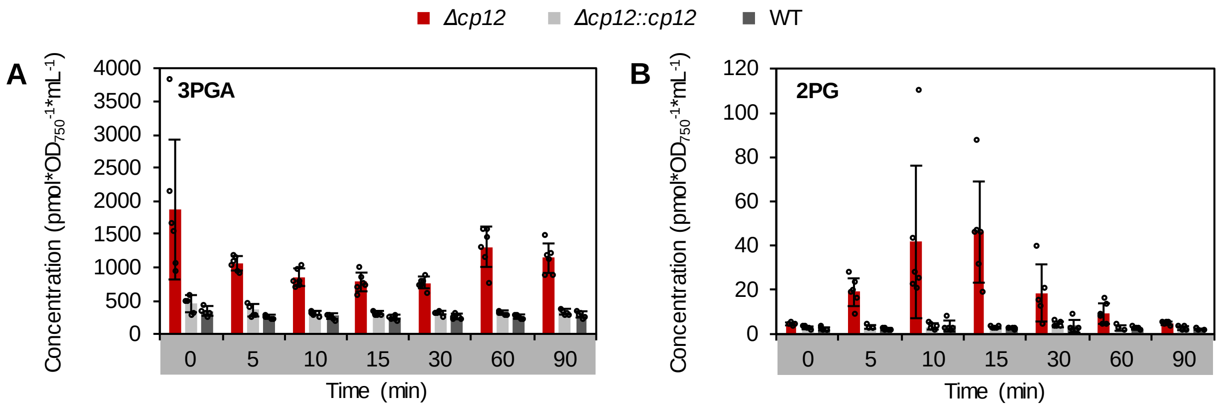

**C** Tukey's HSD

| Time (min) | 0 | 5 | 10 | 15 | 30 | 60 | 90 |
|---|---|---|---|---|---|---|---|
| *Δcp12* | b | c | ac | ac | ac | bc | c |
| *Δcp12::cp12* | a | a | a | a | a | a | a |
| WT | a | a | a | a | a | a | a |

**D** Tukey's HSD

| Time (min) | 0 | 5 | 10 | 15 | 30 | 60 | 90 |
|---|---|---|---|---|---|---|---|
| *Δcp12* | a | ab | bc | c | ab | a | a |
| *Δcp12::cp12* | a | a | a | a | a | a | a |
| WT | a | a | a | a | a | a | a |

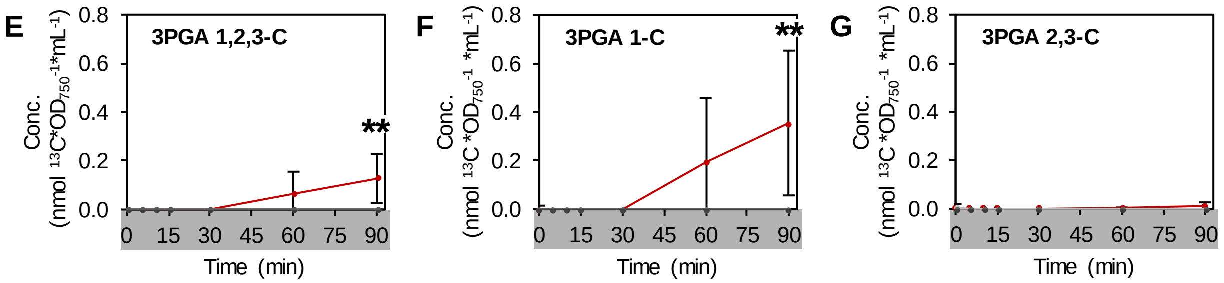

**Figure S 3. Elevated levels of 3PGA and 2PG and minor $CO_2$ assimilation at the end of the night in *Δcp12*.** *Δcp12*, *Δcp12::cp12* and WT were labeled with $^{13}CO_2$ and sampled at the end of the fourth night ($t_0$: 90 min before onset of day settings). Samples were analyzed with GC-MS. 3PGA (**A**) and 2PG (**B**)concentration (pmol*$OD_{750}$*$mL^{-1}$) in *Δcp12* compared to *Δcp12::cp12* and WT. **E-G** E$^{13}$C concnetration (pmol*$OD_{750}$*$mL^{-1}$) of 3PGA 1,2,3-C (E), 1-C (F) and 2,3-C (G). Data points show the average of 5 independent biological replicates ± standard deviation. Asterisks show significant differences between *Δcp12* and WT and *Δcp12::cp12* (tested independently) based on Wilcoxon-Mann-Whitney testing (**: $p<0.01$). Letters in the table (**C+D**) display significant differences of 3PGA concentration between all conditions based on Tukey's HSD ($p<0.05$).

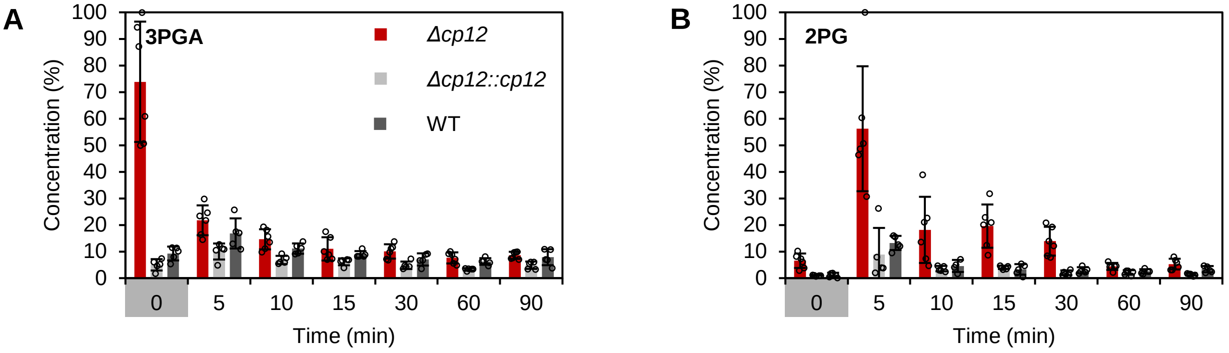

**C** **Tukey's HSD**

| Time (min) | 0 | 5 | 10 | 15 | 30 | 60 | 90 |
|---|---|---|---|---|---|---|---|
| *Δcp12* | c | b | ab | ab | ab | a | a |
| *Δcp12::cp12* | a | ab | a | a | a | a | a |
| WT | ab | ab | ab | ab | a | a | a |

**D** **Tukey's HSD**

| Time (min) | 0 | 5 | 10 | 15 | 30 | 60 | 90 |
|---|---|---|---|---|---|---|---|
| *Δcp12* | abc | d | bc | c | abc | ab | abc |
| *Δcp12::cp12* | a | abc | ab | abc | a | a | a |
| WT | a | abc | abc | ab | ab | ab | ab |

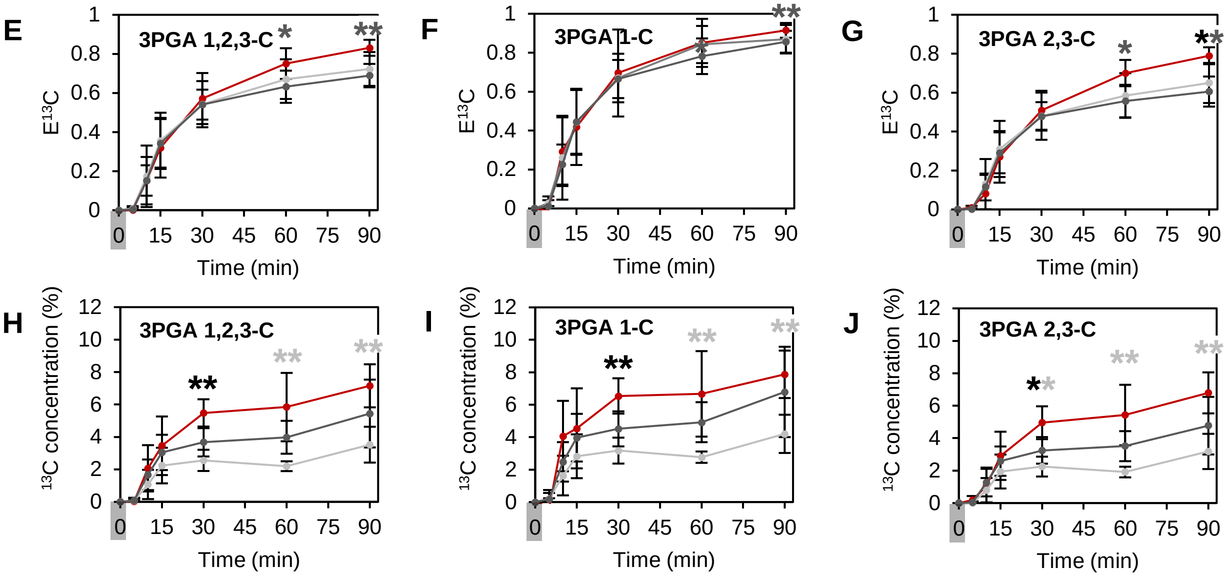

**Figure S 4. $CO_2$ assimilation by RUBISCO at the beginning of the day in *Δcp12*.** *Δcp12*, *Δcp12::cp12* and WT were labeled with $^{13}CO_2$ and sampled at transition from night 4 to the next day ($t_0$: shortly before onset of light, $^{13}CO_2$ directly after light on). Samples were analyzed with GC-MS. Relative 3PGA (**A**) and 2PG (**B**)concentration (%) in *Δcp12* compared to *Δcp12::cp12* and WT. **E-G** Fractional $E^{13}C$ in 1,2,3-C (**E**), 2,3-C (**F**) and 1-C (**G**) of 3PGA. **H-J** $E^{13}C$ concentration (%) in 1,2,3-C (**H**), 2,3-C (**I**) and 1-C (**J**) of 3PGA. Data points show the average of 5 independent biological replicates ± standard deviation. Asterisks show significant differences between *Δcp12* and WT and *Δcp12::cp12* (tested independently) based on Wilcoxon-Mann-Whitney testing (**: $p<0.01$). Letters in the table (**C+D**) display significant differences of 3PGA concentration between all conditions based on Tukey's HSD ($p<0.05$).

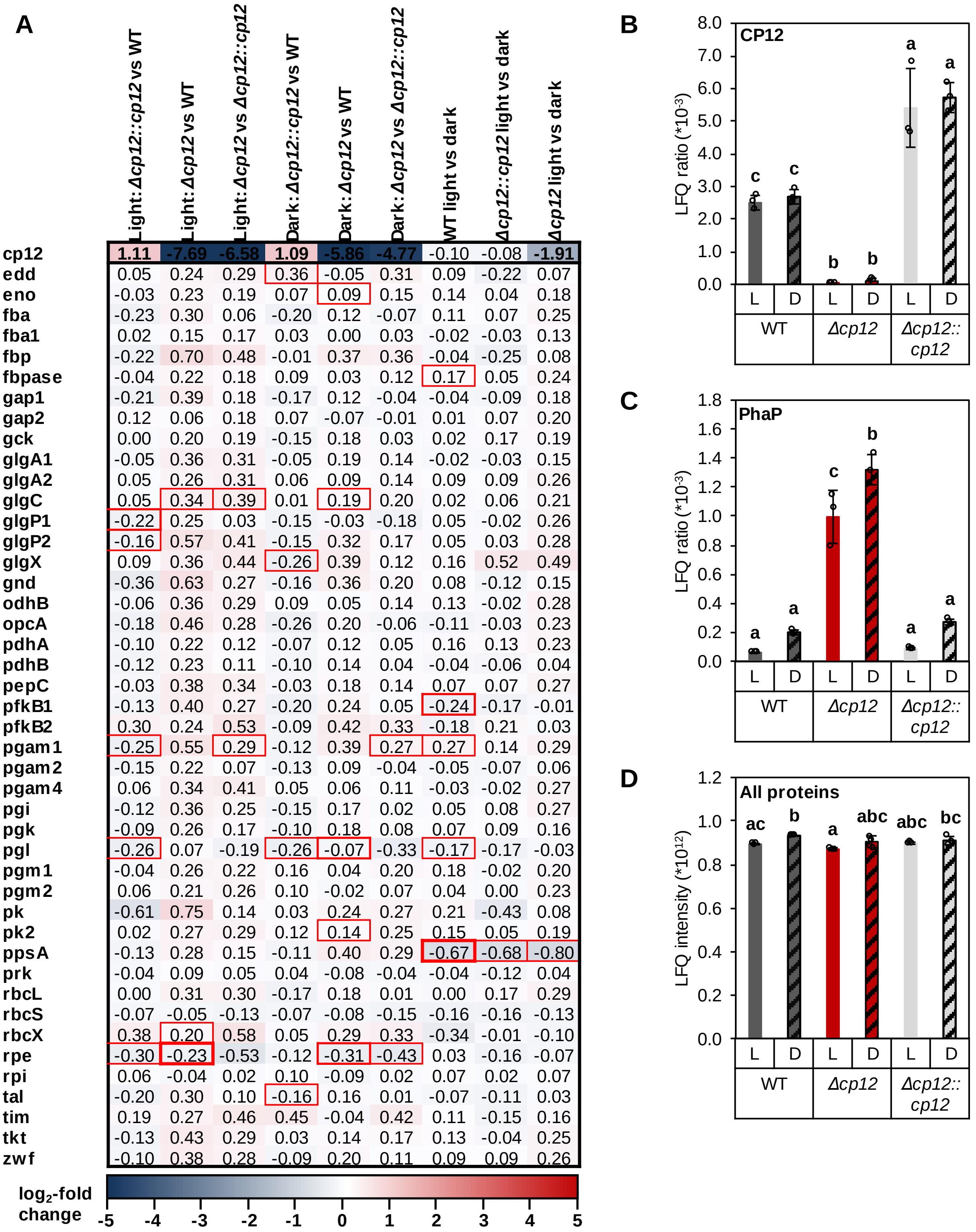

| | Light: Δcp12::cp12 vs WT | Light: Δcp12 vs WT | Light: Δcp12 vs Δcp12::cp12 | Dark: Δcp12::cp12 vs WT | Dark: Δcp12 vs WT | Dark: Δcp12 vs Δcp12::cp12 | WT light vs dark | Δcp12::cp12 light vs dark | Δcp12 light vs dark |
|---|---|---|---|---|---|---|---|---|---|
| cp12 | 1.11 | -7.69 | -6.58 | 1.09 | -5.86 | -4.77 | -0.10 | -0.08 | -1.91 |
| edd | 0.05 | 0.24 | 0.29 | 0.36 | -0.05 | 0.31 | 0.09 | -0.22 | 0.07 |
| eno | -0.03 | 0.23 | 0.19 | 0.07 | 0.09 | 0.15 | 0.14 | 0.04 | 0.18 |
| fba | -0.23 | 0.30 | 0.06 | -0.20 | 0.12 | -0.07 | 0.11 | 0.07 | 0.25 |
| fba1 | 0.02 | 0.15 | 0.17 | 0.03 | 0.00 | 0.03 | -0.02 | -0.03 | 0.13 |
| fbp | -0.22 | 0.70 | 0.48 | -0.01 | 0.37 | 0.36 | -0.04 | -0.25 | 0.08 |
| fbpase | -0.04 | 0.22 | 0.18 | 0.09 | 0.03 | 0.12 | 0.17 | 0.05 | 0.24 |
| gap1 | -0.21 | 0.39 | 0.18 | -0.17 | 0.12 | -0.04 | -0.04 | -0.09 | 0.18 |
| gap2 | 0.12 | 0.06 | 0.18 | 0.07 | -0.07 | -0.01 | 0.01 | 0.07 | 0.20 |
| gck | 0.00 | 0.20 | 0.19 | -0.15 | 0.18 | 0.03 | 0.02 | 0.17 | 0.19 |
| glgA1 | -0.05 | 0.36 | 0.31 | -0.05 | 0.19 | 0.14 | -0.02 | -0.03 | 0.15 |
| glgA2 | 0.05 | 0.26 | 0.31 | 0.06 | 0.09 | 0.14 | 0.09 | 0.09 | 0.26 |
| glgC | 0.05 | 0.34 | 0.39 | 0.01 | 0.19 | 0.20 | 0.02 | 0.06 | 0.21 |
| glgP1 | -0.22 | 0.25 | 0.03 | -0.15 | -0.03 | -0.18 | 0.05 | -0.02 | 0.26 |
| glgP2 | -0.16 | 0.57 | 0.41 | -0.15 | 0.32 | 0.17 | 0.05 | 0.03 | 0.28 |
| glgX | 0.09 | 0.36 | 0.44 | -0.26 | 0.39 | 0.12 | 0.16 | 0.52 | 0.49 |
| gnd | -0.36 | 0.63 | 0.27 | -0.16 | 0.36 | 0.20 | 0.08 | -0.12 | 0.15 |
| odhB | -0.06 | 0.36 | 0.29 | 0.09 | 0.05 | 0.14 | 0.13 | -0.02 | 0.28 |
| opcA | -0.18 | 0.46 | 0.28 | -0.26 | 0.20 | -0.06 | -0.11 | -0.03 | 0.23 |
| pdhA | -0.10 | 0.22 | 0.12 | -0.07 | 0.12 | 0.05 | 0.16 | 0.13 | 0.23 |
| pdhB | -0.12 | 0.23 | 0.11 | -0.10 | 0.14 | 0.04 | -0.04 | -0.06 | 0.04 |
| pepC | -0.03 | 0.38 | 0.34 | -0.03 | 0.18 | 0.14 | 0.07 | 0.07 | 0.27 |
| pfkB1 | -0.13 | 0.40 | 0.27 | -0.20 | 0.24 | 0.05 | -0.24 | -0.17 | -0.01 |
| pfkB2 | 0.30 | 0.24 | 0.53 | -0.09 | 0.42 | 0.33 | -0.18 | 0.21 | 0.03 |
| pgam1 | -0.25 | 0.55 | 0.29 | -0.12 | 0.39 | 0.27 | 0.27 | 0.14 | 0.29 |
| pgam2 | -0.15 | 0.22 | 0.07 | -0.13 | 0.09 | -0.04 | -0.05 | -0.07 | 0.06 |
| pgam4 | 0.06 | 0.34 | 0.41 | 0.05 | 0.06 | 0.11 | -0.03 | -0.02 | 0.27 |
| pgi | -0.12 | 0.36 | 0.25 | -0.15 | 0.17 | 0.02 | 0.05 | 0.08 | 0.27 |
| pgk | -0.09 | 0.26 | 0.17 | -0.10 | 0.18 | 0.08 | 0.07 | 0.09 | 0.16 |
| pgl | -0.26 | 0.07 | -0.19 | -0.26 | -0.07 | -0.33 | -0.17 | -0.17 | -0.03 |
| pgm1 | -0.04 | 0.26 | 0.22 | 0.16 | 0.04 | 0.20 | 0.18 | -0.02 | 0.20 |
| pgm2 | 0.06 | 0.21 | 0.26 | 0.10 | -0.02 | 0.07 | 0.04 | 0.00 | 0.23 |
| pk | -0.61 | 0.75 | 0.14 | 0.03 | 0.24 | 0.27 | 0.21 | -0.43 | 0.08 |
| pk2 | 0.02 | 0.27 | 0.29 | 0.12 | 0.14 | 0.25 | 0.15 | 0.05 | 0.19 |
| ppsA | -0.13 | 0.28 | 0.15 | -0.11 | 0.40 | 0.29 | -0.67 | -0.68 | -0.80 |
| prk | -0.04 | 0.09 | 0.05 | 0.04 | -0.08 | -0.04 | -0.04 | -0.12 | 0.04 |
| rbcL | 0.00 | 0.31 | 0.30 | -0.17 | 0.18 | 0.01 | 0.00 | 0.17 | 0.29 |
| rbcS | -0.07 | -0.05 | -0.13 | -0.07 | -0.08 | -0.15 | -0.16 | -0.16 | -0.13 |
| rbcX | 0.38 | 0.20 | 0.58 | 0.05 | 0.29 | 0.33 | -0.34 | -0.01 | -0.10 |
| rpe | -0.30 | -0.23 | -0.53 | -0.12 | -0.31 | -0.43 | 0.03 | -0.16 | -0.07 |
| rpi | 0.06 | -0.04 | 0.02 | 0.10 | -0.09 | 0.02 | 0.07 | 0.02 | 0.07 |
| tal | -0.20 | 0.30 | 0.10 | -0.16 | 0.16 | 0.01 | -0.07 | -0.11 | 0.03 |
| tim | 0.19 | 0.27 | 0.46 | 0.45 | -0.04 | 0.42 | 0.11 | -0.15 | 0.16 |
| tkt | -0.13 | 0.43 | 0.29 | 0.03 | 0.14 | 0.17 | 0.13 | -0.04 | 0.25 |
| zwf | -0.10 | 0.38 | 0.28 | -0.09 | 0.20 | 0.11 | 0.09 | 0.09 | 0.26 |

**Figure S 5. Proteomics analysis of *Δcp12*, *Δcp12::cp12* and WT at the end of the day and during the night (90 min darkness). A** $Log_2$-fold changes were calculated based on LFQ ratios (LFQ intensities normalized by sum of LFQ intensities per sample) between strains and conditions for all proteins known to be involved in central carbon metabolism. Significant changes are surrounded by red lines (line width depends on p-value based on student's t-test: $p<0.05$: 0.5 pt, $p<0.01$: 1 pt, $p<0.001$: 1.5 pt). **B** Protein abundance (LFQ ratio) is displayed for CP12 to proof the function of *Δcp12*. **C** PhaP, involved in regulating the size of PhB granula and potentially PHB synthase activity, strongly accumulates in *Δcp12* during day and night conditions. **D** Sum of all protein LFQ intensities per sample in the different conditions which was used for the normalization of LFQ intensities. Letters indicate significance based on Tukey's HSD ($p<0.05$).

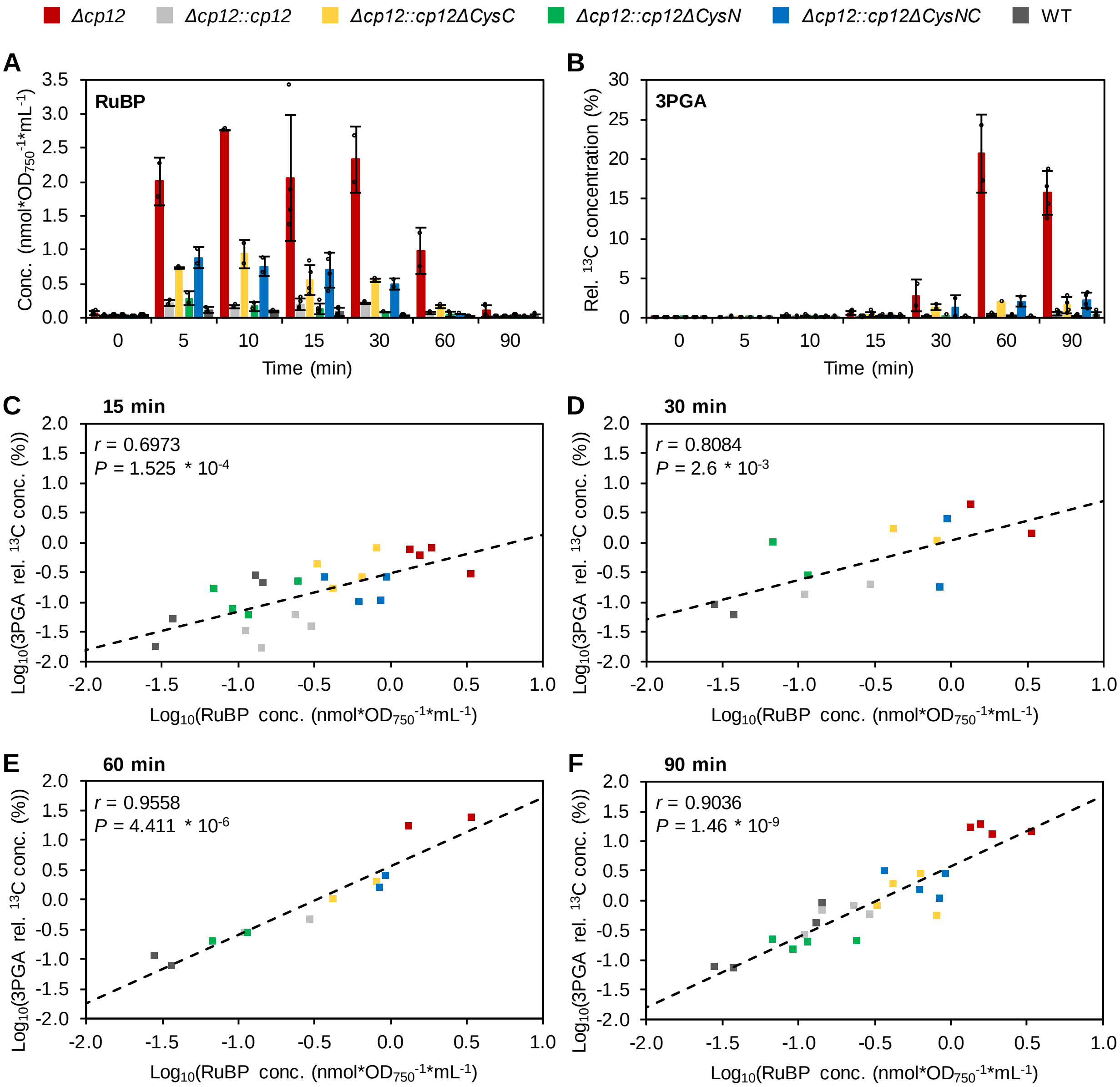

**Figure S 6. Time-shifted correlation of RuBP concentration and 3PGA relative $^{13}C$ concentration.** *Synechocystis* WT and mutant strains were labeled and sampled at transition to the night. Metabolites were quantified and $^{13}C$ enrichment analyzed with LC-MS/MS. **A** RuBP accumulates in *Δcp12* directly after onset of the night, but also to a smaller amount in *Δcp12::cp12ΔCysC* and *Δcp12::cp12ΔCysNC*. The RuBP level decrease after 60 min in all strains. **B** The relative $^{13}C$ concentration (max-scaled rel. concentration multiplied with $^{13}C$ enrichment (%)) of 3PGA increases after 15 to 90 min especially in *Δcp12*, but also in *Δcp12::cp12ΔCysC* and *Δcp12::cp12ΔCysNC*. We correlated RuBP concentration at 15 min with 3PGA relative $^{13}C$ concentration at 15 min (**C**), 30 min (**D**), 60 min (**E**) and 90 min (**F**). The correlation was strongest at 60 and 90 min with $r > 0.9$ and highly significant with $P < 5*10^{-6}$.

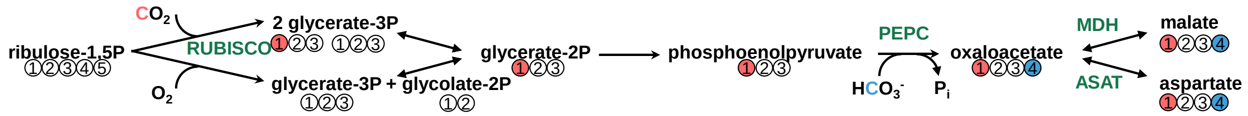

**Figure S 7. Carbon organization during RUBISCO reaction, glycolysis and PEPC reaction.** Carboxylation reaction of RUBISCO leads to 2 3PGA molecules. The newly assimilated $C_i$ will be found in one of the 3PGA molecules in 1-C. The organization of the carbon backbone is kept from 3PGA over 2-Phosphoglycerate (2PGA) to PEP. PEPC adds $HCO_3$- to PEP resulting in oxaloacetate having the newly assimilated $C_i$ at 4-C. This is transferred to malate and aspartate and carbon organization is kept. Thus, 1-C of aspartate gives information about RUBISCO activity, 4-C about PEPC activity and 2,3-C about regeneration of RuBP. Enzyme abbreviations: MDH – Malate dehydrogenase, ASAT – Aspartate aminotransferase.
